# Development of potency, breadth and resilience to viral escape mutations in SARS-CoV-2 neutralizing antibodies

**DOI:** 10.1101/2021.03.07.434227

**Authors:** Frauke Muecksch, Yiska Weisblum, Christopher O. Barnes, Fabian Schmidt, Dennis Schaefer-Babajew, Julio C C Lorenzi, Andrew I Flyak, Andrew T DeLaitsch, Kathryn E Huey-Tubman, Shurong Hou, Celia A. Schiffer, Christian Gaebler, Zijun Wang, Justin Da Silva, Daniel Poston, Shlomo Finkin, Alice Cho, Melissa Cipolla, Thiago Y. Oliveira, Katrina G. Millard, Victor Ramos, Anna Gazumyan, Magdalena Rutkowska, Marina Caskey, Michel C. Nussenzweig, Pamela J. Bjorkman, Theodora Hatziioannou, Paul D. Bieniasz

## Abstract

Antibodies elicited in response to infection undergo somatic mutation in germinal centers that can result in higher affinity for the cognate antigen. To determine the effects of somatic mutation on the properties of SARS-CoV-2 spike receptor-binding domain (RBD)-specific antibodies, we analyzed six independent antibody lineages. As well as increased neutralization potency, antibody evolution changed pathways for acquisition of resistance and, in some cases, restricted the range of neutralization escape options. For some antibodies, maturation apparently imposed a requirement for multiple spike mutations to enable escape. For certain antibody lineages, maturation enabled neutralization of circulating SARS-CoV-2 variants of concern and heterologous sarbecoviruses. Antibody-antigen structures revealed that these properties resulted from substitutions that allowed additional variability at the interface with the RBD. These findings suggest that increasing antibody diversity through prolonged or repeated antigen exposure may improve protection against diversifying SARS-CoV-2 populations, and perhaps against other pandemic threat coronaviruses.

## INTRODUCTION

Neutralizing antibodies elicited by infection or vaccination are a central component of immunity to subsequent challenge by viruses (Plotkin, 2010) and can also confer passive immunity in prophylactic or therapeutic settings. In the case of SARS-CoV-2, an understanding of how viral variants evade antibodies, and how affinity maturation might generate antibodies that are resilient to viral evolution is important to guide vaccination and treatment strategies.

The receptor-binding domains (RBDs) of SARS-CoV-2 spike trimer are key neutralization targets and potent RBD-specific antibodies have been isolated from many convalescent donors (Brouwer et al., 2020; Cao et al., 2020; Chen et al., 2020; Chi et al., 2020; Hansen et al., 2020; Ju et al., 2020; Kreer et al., 2020; Robbiani et al., 2020; Rogers et al., 2020; Seydoux et al., 2020; Shi et al., 2020; Wec et al., 2020; Wu et al., 2020b; Zost et al., 2020). Indeed, such antibodies are used for treatment of SARS-CoV-2 infection (Chen et al., 2021; Weinreich et al., 2021). Typically, RBD-specific neutralizing antibodies isolated during early convalescence have low levels of somatic hypermutation and nearly identical antibodies derived from specific rearranged antibody genes (e.g., *VH3-53*/*VH3-63*) (Barnes et al., 2020c; Robbiani et al., 2020; Yuan et al., 2020) are found in distinct convalescent or vaccinated individuals (Wang et al., 2021). Consistent with these findings, high titer neutralizing sera are generated following administration of at least some SARS-CoV-2 vaccines (Sahin et al., 2020; Widge et al., 2021). Conversely, SARS-CoV-2 infection may sometimes fail to induce sufficient B-cell stimulation and expansion to generate high neutralizing antibody titers. Indeed, neutralizing titers are low in some convalescent individuals, including those from whom commonly elicited potent antibodies can be cloned (Luchsinger et al., 2020; Robbiani et al., 2020; Wu et al., 2020a).

The RBD exhibits flexibility and binds the ACE2 receptor only in an “up” conformation, not in the “down” RBD conformation of the closed, prefusion trimer (Walls et al., 2020; Wrapp et al., 2020). Structural studies have allowed designation of distinct RBD-binding antibody structural classes (Barnes et al., 2020b). Class 1 antibodies are derived from *VH3-53* or *VH3-63* gene segments, include short CDRH3s, and recognize the ACE2 binding site on RBDs in an “up” conformation (Barnes et al., 2020b; Barnes et al., 2020c; Hurlburt et al., 2020; Shi et al., 2020; Wu et al., 2020c; Yuan et al., 2020). Class 2 antibodies are derived from a variety of VH gene segments, also target the ACE2 binding site, but can bind to RBDs in either an “up” or “down” conformation. Some class 2 antibodies, e.g., C144 and S2M11 (Barnes et al., 2020b; Tortorici et al., 2020), bridge adjacent “down” RBDs to lock the spike trimer into a closed prefusion conformation. Class 3 antibodies, which can recognize “up” or “down” RBDs, do not target the ACE2 binding site (Barnes et al., 2020b).

Despite the fact that cloned RBD-specific antibodies can select resistance mutations, such as E484K, in cell culture (Baum et al., 2020; Weisblum et al., 2020), until recently, little evidence had emerged that antibodies have imposed selective pressure on circulating SARS-CoV-2 populations. Nevertheless, variability and decay of convalescent neutralizing titers, (Gaebler et al., 2021; Luchsinger et al., 2020; Muecksch et al., 2020; Robbiani et al., 2020; Seow et al., 2020), suggests that reinfection by SARS-CoV-2 may occur at some frequency. Indeed, recent reports have documented reinfection or rapidly increasing case numbers, associated with SARS-CoV-2 variants with resistance to commonly elicited antibodies (Fujino et al., 2021; Tegally et al., 2020; Volz et al., 2021; Wang et al., 2021; West et al., 2021; Wibmer et al., 2021).

The majority of SARS-CoV-2 antibodies that have been studied in detail were cloned from individuals early in convalescence and have relatively low levels of somatic mutation. However, recent work has shown that antibodies evolve in convalescent patients, accumulating somatic mutations that can affect function (Gaebler et al., 2021; Sakharkar et al., 2021; Sokal et al., 2021). Here, we present a detailed functional and structural characterization of several groups of clonally-related antibodies recovered from the same individuals shortly after infection and then later in convalescence. We show that somatic mutations acquired in the months after infection endow some SARS-CoV-2 RBD-specific antibodies with greater neutralization potency and breadth. Most importantly, the acquisition of somatic mutations provides some antibodies with resilience to viral mutations that would otherwise enable SARS-CoV-2 to escape their neutralizing effects.

## Results

### Evaluating resistance to clonally-related SARS-CoV-2 neutralizing antibodies

To determine the functional consequences of antibody maturation in SARS-CoV-2 convalescence, we compared clonally-related antibodies, from six lineages (Table S1), isolated at a mean of 1.3 or 6.2 months (m) after PCR diagnosis of SARS-CoV-2 infection from patients with mild to moderately severe disease (Gaebler et al., 2021; Robbiani et al., 2020). We measured neutralization potency against a panel of HIV-1 SARS-CoV-2 pseudotypes bearing single amino acid substitutions that are naturally occurring or known to confer resistance to neutralization by individual human antibodies or plasma (Weisblum et al., 2020). Additionally, to determine the ability of the SARS-CoV-2 spike to acquire mutations conferring resistance to the antibodies, we employed a pair of replication-competent chimeric VSV derivatives (rVSV/SARS-CoV-2_1D7 / 2E1_) (Schmidt et al., 2020; Weisblum et al., 2020) to select antibody escape variants.

### Effects of maturation on potency and resilience of class 2 SARS-CoV-2 neutralizing antibodies

Class 2 anti-RBD antibodies are commonly elicited and recognize an epitope that includes E484 (Barnes et al., 2020b), a site that is mutated in certain circulating SARS-CoV-2 ‘variants of concern’ (Fujino et al., 2021; West et al., 2021; Wibmer et al., 2021). We examined members of three class 2 antibody lineages.

#### The C144/C051/C052 lineage

One group of class 2, *VH3-53/VL2-14*-encoded antibodies included C144, a potent neutralizing antibody (IC_50_ <10ng/ml) isolated at 1.3m of convalescence (Robbiani et al., 2020) that is in clinical development for therapy/prophylaxis. Two clonally-related antibodies isolated from the C144 donor at 6.2m included C051, which was marginally less potent (IC_50_ ∼25ng/ml), and C052, which had similar potency to C144 (Gaebler et al., 2021).

SARS-CoV-2 pseudotype neutralization assay revealed that several RBD substitutions at positions L455, F456, E484, F490, Q493 and S494, which conferred C144 resistance (Figure1A, B), had no effect C051 and C052 sensitivity. Some, but not all, naturally occurring E484 substitutions that conferred C144 resistance also conferred resistance to C051 and/or C052. Selection for rVSV/SARS-CoV-2 resistance mutations using C144 gave enrichment of multiple substitutions at two positions; E484K/A/G and Q493R/K (Figure 1C), and plaque purification from selected virus populations yielded isolates with E484K or Q493R substitutions that confer high level C144 resistance (Weisblum et al., 2020) (Figure S1A). Conversely, rVSV/SARS-CoV-2 replication in the presence of C051 and C052 led to the dominance of the E484K mutant only, and rVSV/SARS-CoV-2 isolates bearing E484K substitutions were resistant to C051 and C052 (Figure 1C, S1). Thus, in this lineage of potently neutralizing antibodies elicited early after infection somatic mutation conferred resilience to a subset of naturally occurring potential escape variants. Nevertheless, resistance to each of the antibodies in this commonly elicited lineage was conferred by E484K (Figure 1A,B, Figure S1).

**Figure 1.**
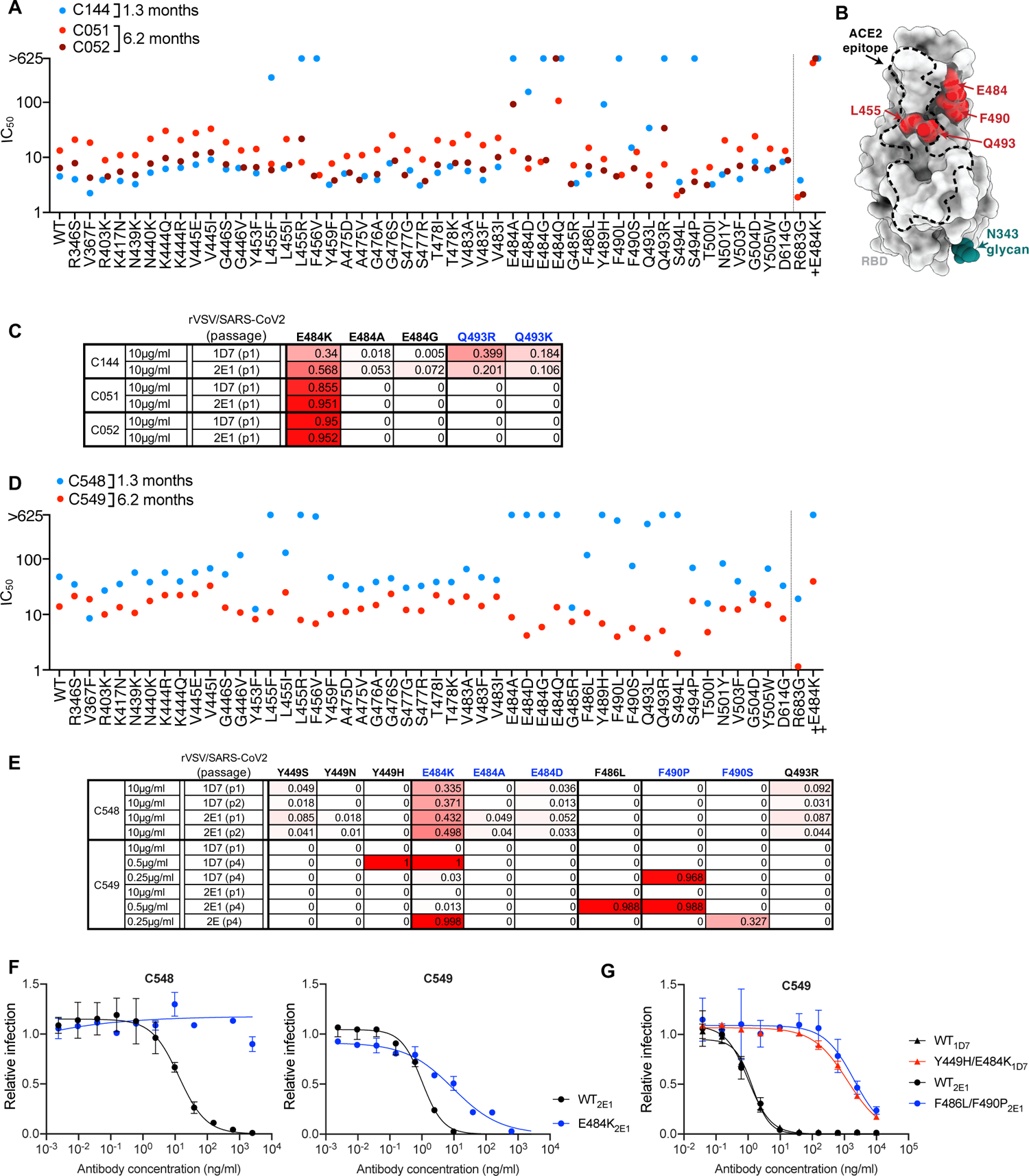
Effects of somatic mutation of class 2 antibodies on potency and viral escape. (A) Neutralization potency (IC_50_) of C144, C051 and C052 measured using HIV-1-based SARS-CoV-2 variant pseudotypes and HT1080/ACE2cl.14 cells. The E484K substitution was constructed in an R683G (furin cleavage site mutant) background to increase infectivity. Mean of two independent experiments. (B) RBD structure indicating positions of substitutions that affect sensitivity to neutralization by class 2 and C144/C05/C052, C143/C164/C055 and C548/549 lineage antibodies. (C) Decimal fraction (color gradient; white = 0, red = 1) of Illumina sequence reads encoding the indicated RBD substitutions following rVSV/SARS-CoV-2 replication (1D7 and 2E1 virus isolates) in the presence of the indicated amounts of antibodies for the indicated number of passages. (D) As in A for antibodies C548 and C549. (E) As in C for antibodies C548 and C549. Reduced antibody concentrations were required for C549 escape. (F,G) C548 (F) and C549 (G) neutralization of rVSV/SARS-CoV-2 1D7, 2E1 or plaque purified mutants thereof isolated following antibody selection, in 293T/ACE2cl.22 cells. Infected (%GFP+) cells relative to no antibody controls, mean and range of two independent experiments is plotted. See also Figure S1, S2, S3

#### The C143/C164/C055 lineage

A second clonally-related antibody lineage, encoded by *VH3-66/VL2-33*, included C143 and C164, isolated at 1.3m and C055, isolated at 6.2m, were from the same individual as the C144 lineage (Gaebler et al., 2021; Robbiani et al., 2020). C143 and C164 had weak neutralizing activity (IC_50_ values ∼300ng/ml to >625ng/ml) against the HIV-1 pseudotype panel (Figure S2A). Conversely, C055 potently neutralized the majority of SARS-CoV-2 spike variant pseudotypes (IC_50_ values of ∼10ng/ml) (Figure S2A). Naturally occurring spike substitutions (at positions A475, T478, E484, G485, and F486) caused loss of C055 potency (Figure S2A), indicating a target epitope close to that of the C144/C051/C052 lineage. Despite their modest potency, rVSV/SARS-CoV-2 replication with C143 or C164 led to the enrichment of T478K/R mutations, and a plaque purified T478R mutant isolate exhibited near-complete resistance to C143 and C164 while an E484K mutant exhibited partial resistance (Figure S2B, C). Conversely rVSV/SARS-CoV-2 selection with C055 yielded G485S/D and F486V/S, substitutions and isolates with G485S, F486S and F486V mutations exhibited near complete resistance to C055 (Figure S2B, D). Overall, maturation of this antibody lineage yielded both greater potency and a change in the selected spike substitutions that yielded neutralization escape (Figure S2B).

#### The C548/C549 lineage

A third class 2 antibody lineage, encoded by *VH1-69/VL9-49*, included C548, isolated at 1.3m, and C549, isolated at 6.2m (Gaebler et al., 2021; Robbiani et al., 2020). C548 was somewhat less potent (IC_50_ ∼50ng/ml) than C549 (IC_50_ ∼15ng/ml). Similar to C144, naturally occurring substitutions at positions L455, F456, E484, S494, Y489, Q493 and S494 caused near complete loss of C548 potency (Figure 1D). Remarkably, however, C549 potency was unaffected or only marginally affected by most of these mutations. E484K conferred partial (∼50-100-fold) resistance (Figure 1D). Selection experiments with C548 led to rapid enrichment of E484K and Q493K C548-resistant rVSV/SARS-CoV-2mutants (Figure 1E, F), consistent with the finding that E484K or Q493R substitutions conferred C548 resistance in the HIV-1 pseudotype assay. In contrast, initial attempts to select C549-resistant rVSV/SARS-CoV-2 mutants failed (Figure 1E). However, by reducing the selecting concentration of C549 and sequential passaging with antibody four times, we obtained rVSV/SARS-CoV-2 populations in which Y449H, E484K, F486L and F490P/S mutations were enriched (Figure 1E). Notably, these selected populations yielded only isolates that encoded two RBD substitutions; one Y449H/E484K and the other F486L/F490P. These viruses exhibited greater (1000-fold), C549-resistance than the E484K single mutant (Figure 1G, Fig S3). Because individual substitutions at E484, F486, and F490 caused only partial or no C549 resistance (Fig 1D, G Figure S3), these data suggest that at least two substitutions are required to confer high-level resistance to C549. Thus, for this pair of antibodies, antibody maturation appeared to heighten the genetic barrier for the acquisition of antibody resistance.

### Maturation confers potency and resilience to escape in a class 1 antibody lineage

A fourth antibody lineage, encoded by *VH3-53/VK3-20* genes, also exhibited striking disparity in activity and breadth when 1.3m and 6.2m clonal relatives were compared. Specifically C098, isolated at 1.3m, displayed minimal activity (IC_50_ > 1000ng/ml) against most SARS-CoV-2 pseudotypes while a clonal relative isolated at 6.2m, C099 had IC_50_ values ranging from ∼15-48 ng/ml for all variants except L455R, for which the IC_50_ was increased to 123ng/ml (Figure 2A, B). In rVSV/SARS-CoV-2 selection experiments, the low potency of C098 was reflected in the modest enrichment of mutations. Nevertheless, there was some enrichment of N460 substitutions (Figure 2B, C), and after two passages, rVSV/SARS-CoV-2 (N460Y) mutant was isolated which displayed nearly complete C098 resistance (Figure 2D).

**Figure 2.**
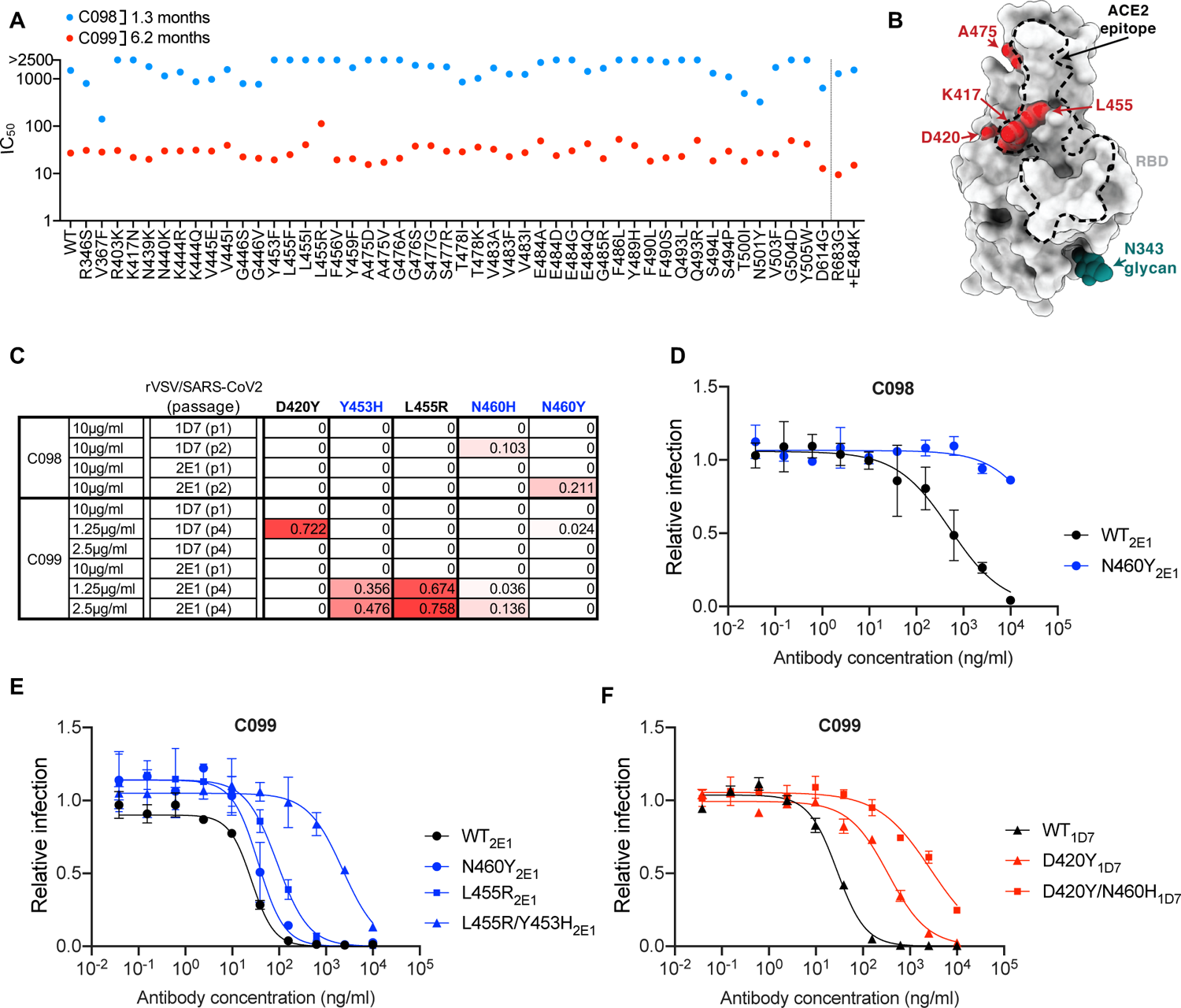
Somatic mutation in a class 1 antibody lineage confers potency and resilience to viral escape. (A) Neutralization potency (IC_50_) of C098 and C099 measured using HIV-1-based SARS-CoV-2 variant pseudotypes and HT1080/ACE2cl.14 cells. The E484K substitution was constructed in an R683G (furin cleavage site mutant) background to increase infectivity. Mean of two independent experiments. (B) RBD structure indicating positions of substitutions that affect sensitivity to neutralization by class 1 and C098 and C099 lineage antibodies. (C) Decimal fraction (color gradient; white = 0, red = 1) of Illumina sequence reads encoding the indicated RBD substitutions following rVSV/SARS-CoV-2 replication (1D7 and 2E1 virus isolates) in the presence of the indicated amounts of antibodies for the indicated number of passages. (D, E, F) C098 (D) and C099 (E, F) neutralization of rVSV/SARS-CoV-2 1D7, 2E1 or plaque purified mutants thereof, isolated following antibody selection, in 293T/ACE2cl.22 cells. Infected (%GFP+) cells relative to no antibody controls, mean and range of two independent experiments is plotted. See also Figure S4

Initial attempts to isolate C099-resistant rVSV/SARS-CoV-2 mutants failed. However, passaging rVSV/SARS-CoV-2 four times with reduced C099 concentrations, yielded populations enriched most prominently in D420Y, Y453H, and L455R substitutions (Figure 2 B, C). Plaque purification yielded D420Y, N460Y, or L455R single mutants with partial (10-fold or less) C099 resistance (Figure 2B, E F) as well as D420Y/N460H and L455R/Y453H double mutants with higher levels of C099-resistance (∼100-fold, Figure 2E, F). Analysis of HIV-1 pseudotypes with these mutations confirmed that D420Y, N460H, L455R and Y453H alone each abolished the weak C098 neutralization activity but conferred no or partial C099-resistance to (Figure S4A, B). However, the D420Y/N460H or L455R/Y453H combinations conferred greater C099-resistance (Figure S4B). Overall, maturation in the C098/99 lineage conferred both potency and resilience to individual spike mutations and appeared to impose a requirement for 2 or more mutations for high-level resistance.

### Acquisition of potency and resilience to escape in class 3 antibody lineages

We next analyzed two pairs of class 3 antibodies, which do not directly compete for ACE2 binding to the SARS-CoV-2 RBD (Barnes et al., 2020b; Weisblum et al., 2020), yet exhibit potent neutralizing activity. Some antibodies in this class, while having very low IC_50_ values, also exhibit incomplete neutralization in pseudotype assays; i.e., a ‘non-neutralizable’ fraction.

#### C132 and C512

The *VH4-4/VL2-14* encoded C132 antibody, isolated at 1.3m, had weak neutralizing activity against the spike variants in the HIV-1 pseudotype assay while its 6.2m clonal derivative, C512, was quite potent, (IC_50_ ∼100ng/ml, Figure 3A). Substitutions at R346, K444 and G446 conferred C512 resistance (Figure 3A, B). Despite its poor potency, rVSV/SARS-2/EGFP replication with C132 generated viral populations enriched for R346 substitutions, and a plaque purified rVSV/SARS-2/EGFP (R346K) mutant that was resistant to the weak activity of C132 (Fig 3B, C, D). In contrast, and despite the fact that R346, K444 and G446 substitutions all conferred resistance in the HIV-1 assay, C512 selected resistant rVSV/SARS-2/EGFP variants with K444 substitutions only (Figure 3C, D). Thus, maturation of the C132/C512 antibody lineage yielded a marked increase in potency, and a concurrent change in the selected resistance mutations.

**Figure 3.**
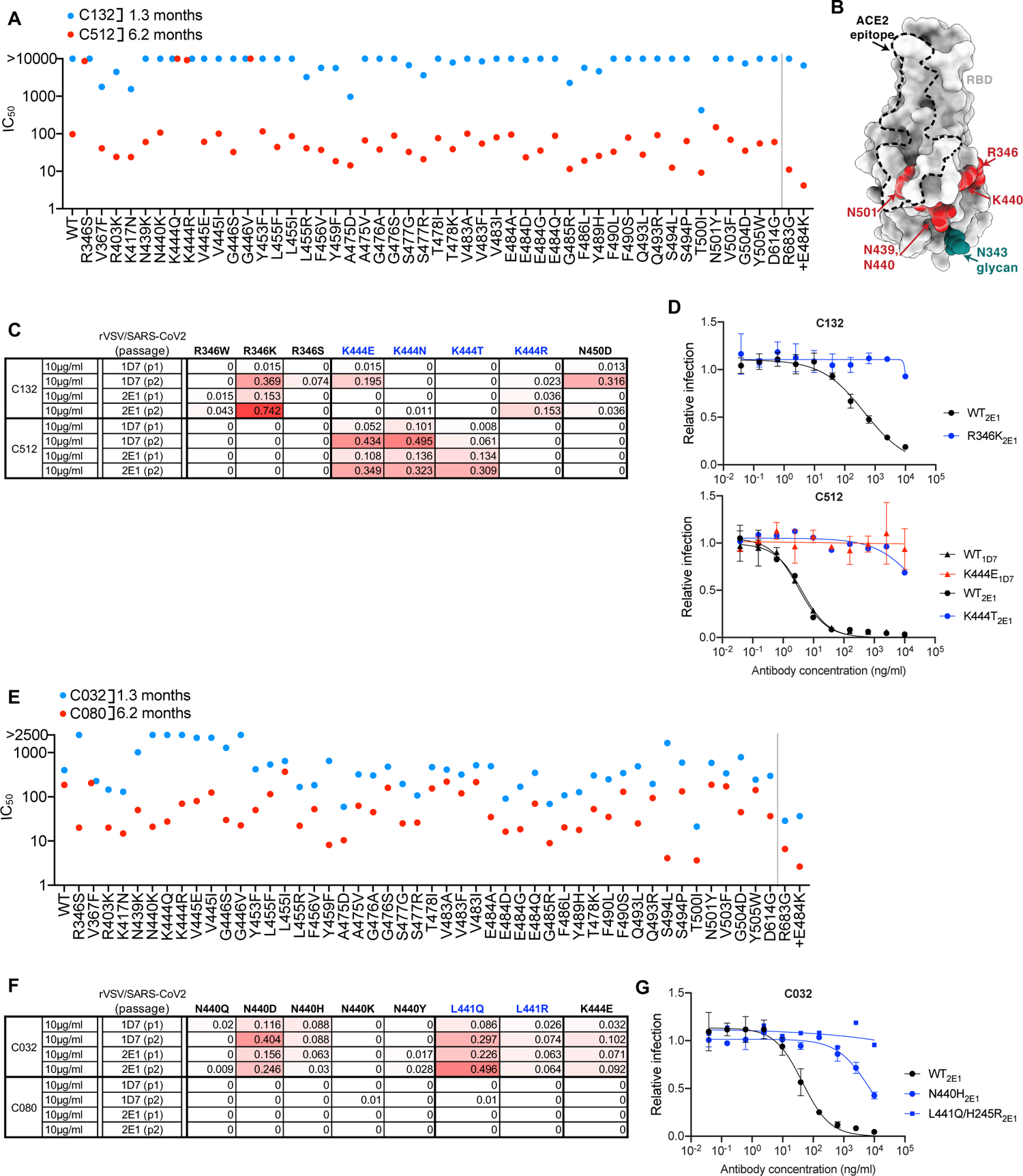
Effects of somatic mutation of class 3 antibodies on potency and viral escape. (A) Neutralization potency (IC_50_) of C132 and C512 measured using HIV-1-based SARS-CoV-2 variant pseudotypes and HT1080/ACE2cl.14 cells. The E484K substitution was constructed in an R683G (furin cleavage site mutant) background to increase infectivity. Mean of two independent experiments. (B) RBD structure indicating positions of substitutions that affect sensitivity to neutralization by class 3 and C132/C512 and C032/C080 lineage antibodies. (C) Decimal fraction (color gradient; white = 0, red = 1) of Illumina sequence reads encoding the indicated RBD substitutions following rVSV/SARS-CoV-2 replication (1D7 and 2E1 virus isolates) in the presence of the indicated amounts of antibodies for the indicated number of passages. (D) C132 and C512 neutralization of rVSV/SARS-CoV-2 1D7, 2E1 or plaque purified mutants thereof, isolated following antibody selection, in 293T/ACE2cl.22 cells. Infected (%GFP+) cells relative to no antibody controls, mean and range of two independent experiments is plotted. (E) As in A for C032 and C080. (F) As in C for C032 and C080. (G) As in D for C032.

#### C032 and C080

For a second clonally-related pair of class 3 antibodies, encoded by *VH5-51/VL1-40*, the antibody isolated at 1.3m (C032) was only ∼2-fold less potent than a derivative isolated at 6.2m (C080). However, mutations at positions R346, N439, N440, K444, V445 and G446 all conferred resistance to C032, but not to C080 (Figure 3B, E). Like some other class 3 antibodies, C032 and C080 exhibited incomplete neutralization complicating selection of rVSV/SARS-CoV-2 resistant variants. Nevertheless, C032 enriched N440 and L441 substitutions, both of which conferred C032 resistance (Figure 3G). Under identical rVSV/SARS-CoV-2 selection conditions, no mutations were enriched in the presence of C080. Therefore, a key property acquired by C080 was resilience to mutations that conferred resistance to its C032 progenitor.

### Neutralizing activity of matured antibodies against RBD ‘variants of concern’

Selection for rVSV/SARS-CoV-2 resistance to class 1, 2 and 3 antibodies in cell culture has repeatedly identified K417, E484 and N501 substitutions, with E484K giving the most pervasive effects against polyclonal plasma (Baum et al., 2020; Greaney et al., 2021; Liu et al., 2021; Wang et al., 2021; Weisblum et al., 2020). We compared the ability of the antibodies studied herein to neutralize pseudotypes with a E484K substitution alone, or in combination with K417N and N501Y substitutions that naturally occur in variants of concern (Fujino et al., 2021; Wibmer et al., 2021) or in combination with L455R, a mutation that affected neutralization by multiple class 1 and class 2 antibodies (Figure 1, Figure 2). The activity of the C144/C051/C052 and C143/C162/C055 class 2 lineages was diminished by the E484K substitution, and there was little scope for additional mutations to further reduce potency (Figure 4A, B). Conversely, while the 1.3m class 2 antibody C548 was inactive against the E484K mutant, its descendent C549 retained partial activity (Figure 4C). C549 activity was modestly further reduced by the K417N/E484K/N501Y combination and abolished by the L455R/E484K combination (Figure 4C), consistent with the notion that 2 substitutions were required to confer maximal C549 escape (Figure 2E, F, Figure S3). The C098/C099 class 1 antibodies were not affected by the E484K mutation or the K417N/E484K/N501Y combination (Figure 4D). The partial loss of potency against the L455R/E484K combination was largely consistent with that seen for the L455R single mutant (Figure S4B). As expected, K417/E484K/N501Y and L455R/E484K mutations did not confer resistance to the class 3 antibodies. In fact, unexpectedly, these mutations sensitized the pseudotypes to neutralization by some class 3 antibodies (Figure 4D). Thus, the E484K substitution, generally undermines the activity of class 2 antibodies, but substitutions found in variants of concern did not impact the activity of the matured class 1 and class 3 antibodies tested herein.

**Figure 4.**
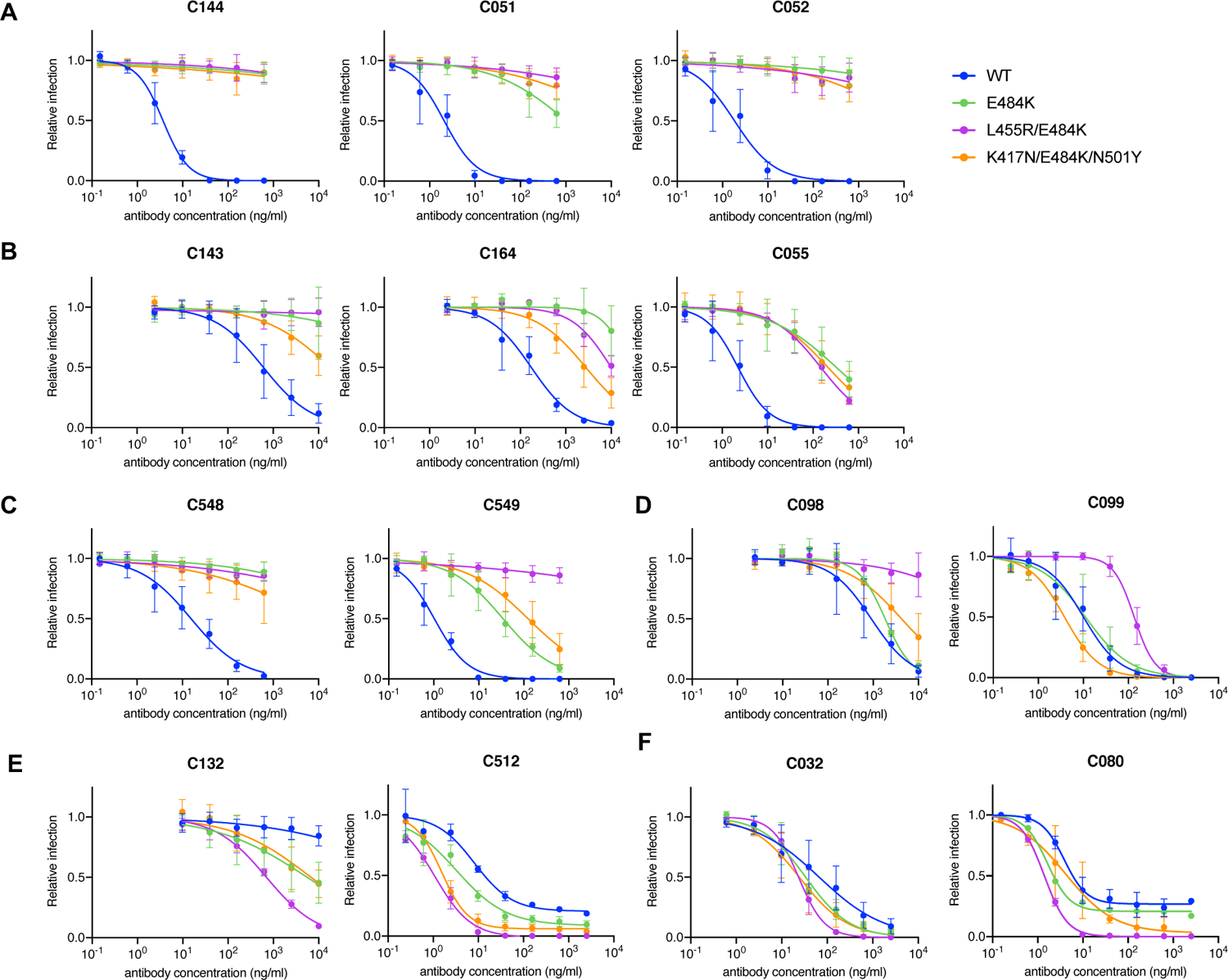
Effect of the E484K substitution alone or in combination with K417N/N501Y or L455R on matured class 1, 2 and 3 antibody sensitivity. (A-F) Neutralization of HIV-1-based SARS-CoV-2 variant pseudotypes by C144/C051/C052 (A), C143/C164/C055 (B), C548/C549 (C), C098/C099 (D), C132/C512 (E) and C032/C080 (F) lineage antibodies in HT1080/ACE2cl.14 cells. Each of these variants was constructed in an R683G (furin cleavage site mutant) background to increase infectivity. Mean and range of two independent experiments.

### Neutralization activity of matured antibodies against sarbecoviruses other than SARS-CoV-2

We next determined whether any of the antibodies could neutralize more divergent sarbecoviruses. SARS-CoV-2 is closely related to the horseshoe bat (*Rinolophus affinis*) coronavirus bCoV-RaTG13 (97.4% amino acid identity in spike) (Zhou et al., 2020), but the SARS-CoV-2 RBD diverges from bCoV-RaTG13 (89.3% identity) and is more closely related (97.4% identity) to pangolin (*Manis javanica*) coronavirus from Guandong, China (pCoV-GD). The RBD of a second pangolin coronavirus found in Guangxi (pCoV-GX) shares 87% amino acid identity with SARS-CoV-2 (Lam et al., 2020; Zhang et al., 2020). The SARS-CoV spike protein is more closely related to coronaviruses found in *Rinolophus Sinicus*, including bCoV-WIV16 with which it shares 94.3% RBD amino acid identity. The SARS-CoV and bat-CoV-WIV16 RBDs share 73-75.4% identity with the SARS-CoV-2 RBD (Li et al., 2005).

None of the antibodies neutralized bCoV-WIV16 pseudotypes (Figure 5A-F). In contrast, all of the antibodies except C512 neutralized pCoV-GD pseudotypes. Some matured antibodies isolated at 6.2m (C055, C549, C099 and C080) neutralized pCoV-GD pseudotypes more potently than their 1.3m clonal predecessors (Figure 5B, C, D, F), recapitulating observations with SARS-CoV-2 pseudotypes. Additionally, C099, unlike its clonally-related predecessor C098, potently neutralized the more distantly related pCoV-GX pseudotype (IC_50_ =16ng/ml, Figure 5D). Finally, the 6.2m class 3 antibody, C080, neutralized SARS-CoV (IC_50_= 71ng/ml). Thus, antibody evolution in SARS-CoV-2 convalescents increased breadth, in some cases enabling neutralization of heterologous pandemic-threat sarbecoviruses.

**Figure 5.**
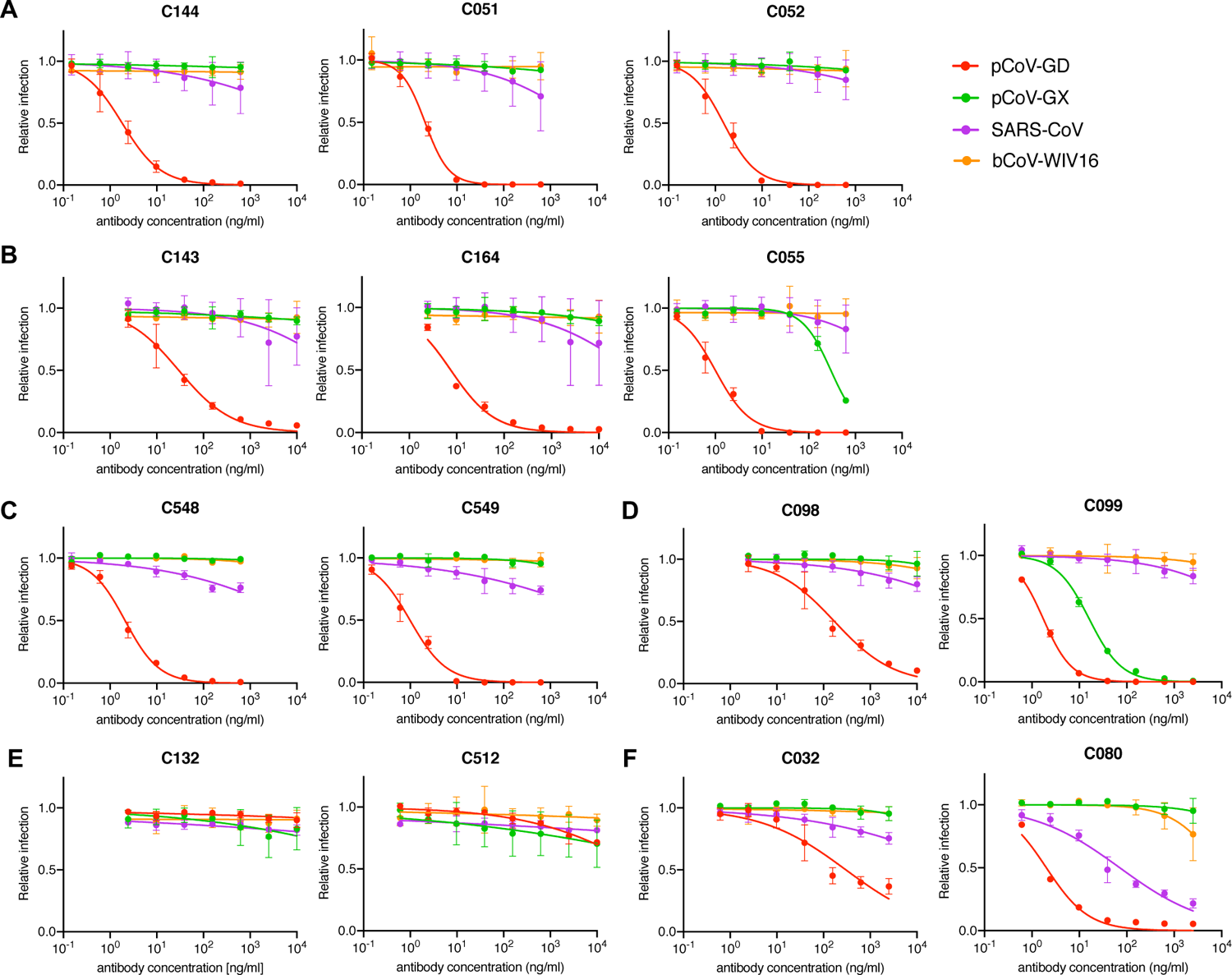
Neutralization of heterologous sarbecoviruses by SARS-CoV-2 elicited antibodies and effects of somatic mutation on breadth. (A-F) Neutralization of HIV-1-based SARS-CoV, bat coronavirus (bCoV WIV16), or pangolin coronaviruses (pCov-GD and pCoV-GX) pseudotypes by C144/C051/C052 (A), C143/C164/C055 (B), C548/C549 (C), C098/C099 (D), C132/C512 (E) and C032/C080 (F) lineage antibodies in HT1080/ACE2cl.14 cells. Mean and range of two independent experiments.

### Structural analyses of somatic hypermutations in antibody clonal pairs

We investigated the effects of somatic mutations on antibody-antigen interactions by solving structures of 1.3m and 6.2m pairs of class 1 (C098/C099) and class 2 (C144/C051) antibody Fab fragments bound to SARS-CoV-2 spike trimers or monomeric RBDs (Figure S5 and Tables S2,3). We also determined structures of 1.3m class 2 (C548) and 1.3m class 3 (C032) Fabs bound to S, allowing modeling of RBD interactions for their 6.2m counterparts (C549 and C080, respectively). Across these structures, most substitutions found at 6.2m post-infection occurred in CDR loops, in or adjacent to antibody paratopes (Figure 6,7, Figure S6,7).

**Figure 6.**
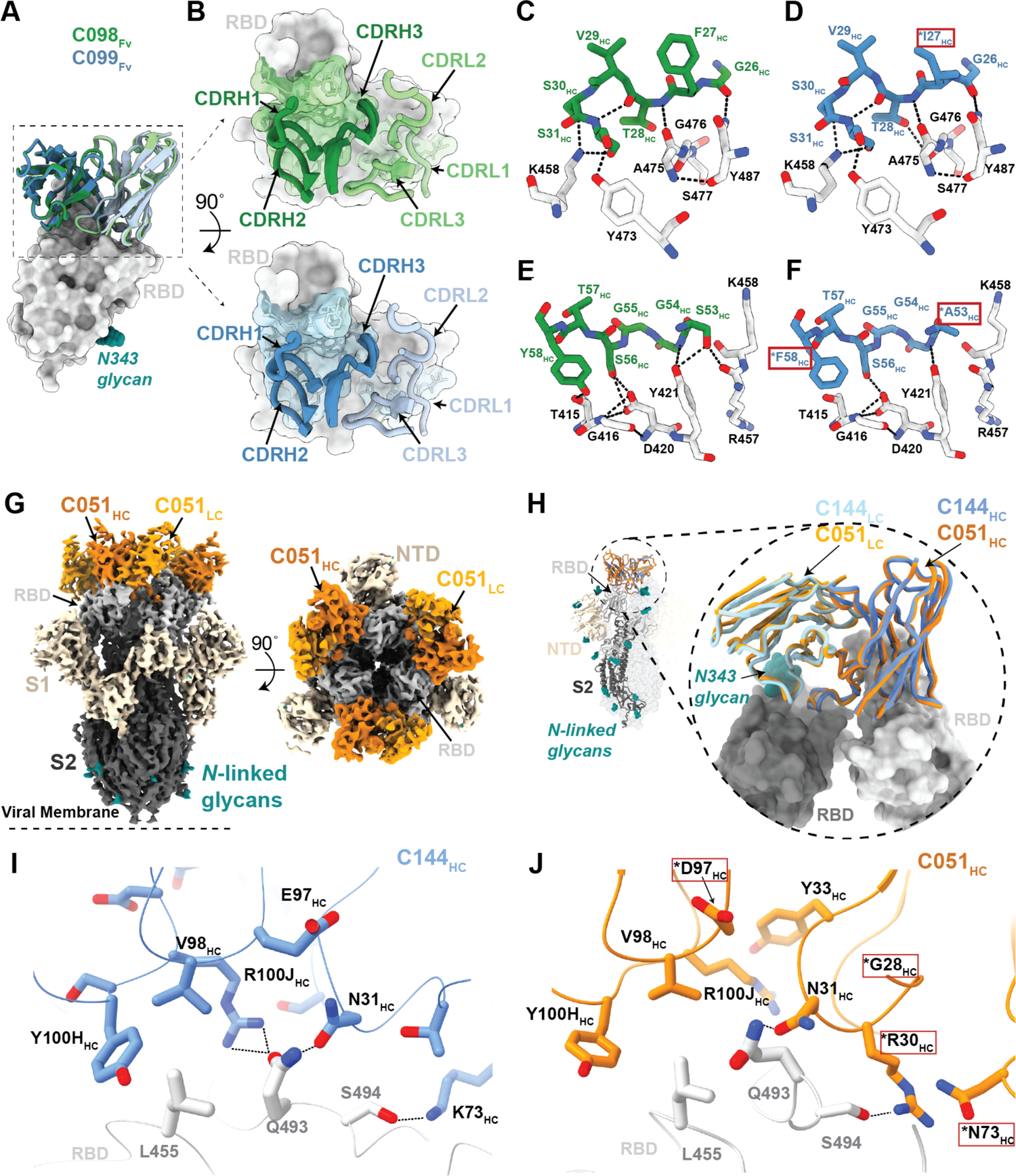
Structures of class 1 and class 2 anti-RBD antibody 1.3m and 6.2m pairs. (A) Overlay ofVwVL domains of class 1 C098 and C099 Fabs bound to RBD from 2.0 A and 2.6 A crystal structures, respectively. (B) CDR loops of C098 and C099 mapped onto the RBD surface. Fab epitopes are colored on the RBD surface. (C,D) Interactions of C098 (panel C) and C099 (panel D) CDRH1 residues with RBD. Residues changed by somatic hypermutation indicated by an asterisk and enclosed in a red box. (E,F) Interactions of C098 (panel E) and C099 (panel F) CDRH2 residues with RBD. Residues changed by somatic hypermutation indicated by an asterisk and enclosed in a red box. (G) 3.5 Å cryo-EM density for class 2 C051-S complex structure (only the V_H_-V_L_ domains of C051 are shown). (H) Overlay of V_H_-V_L_ domains of C051 and C144 Fabs bound to S trimer. Both Fabs bridge between adjacent “down” RBDs, shown in inset as dark and light gray surfaces. (I,J) Interactions between RBD and C144 (panel I) and C051 (panel J) with a subset of interacting residues highlighted as sticks. Potential hydrogen bonds shown as dotted lines. Residues changed by somatic hypermutation indicated by an asterisk and enclosed in a red box. See also Figure S5, S6

**Figure 7.**
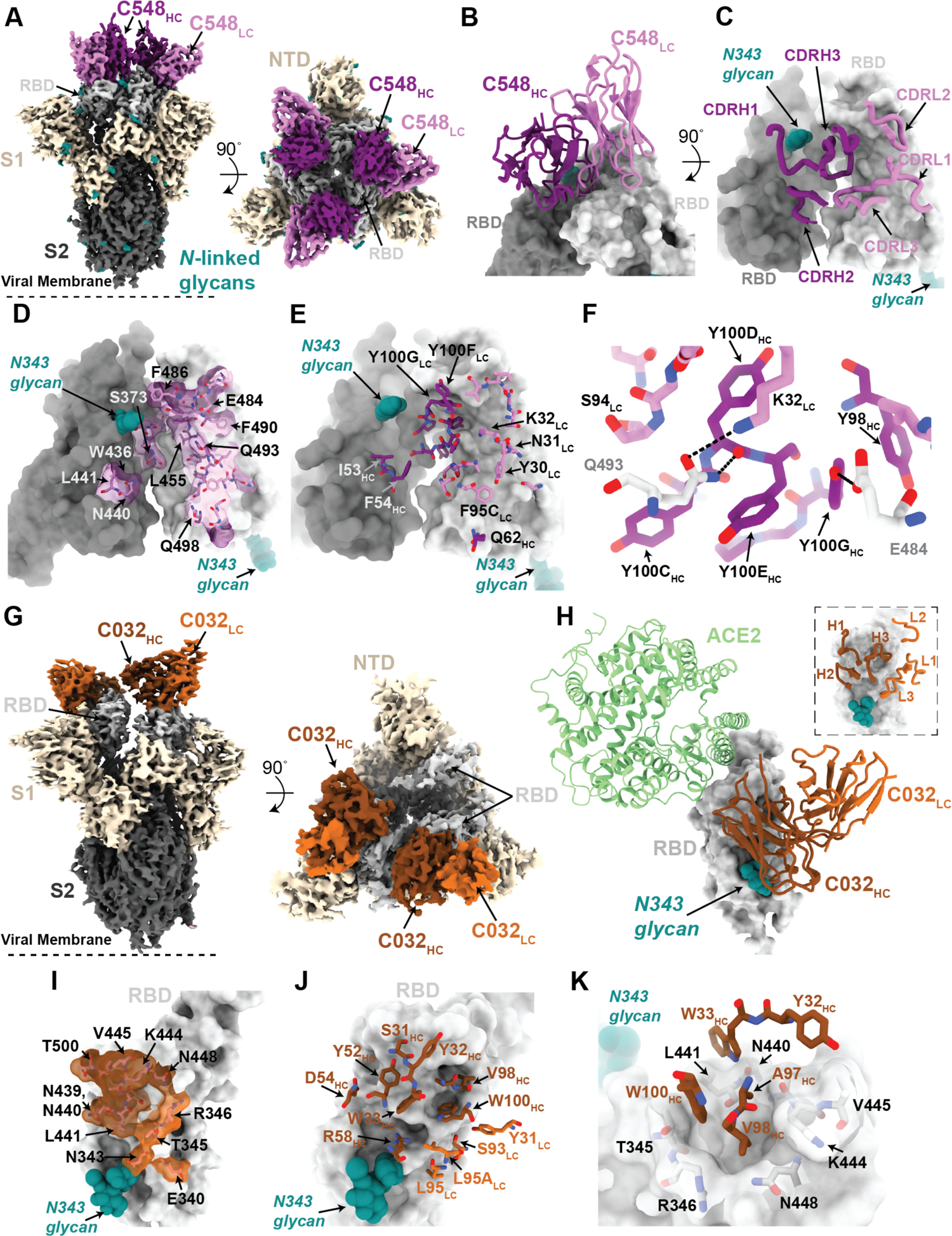
Structures of class 2 and class 3 anti-RBD 1.3m antibodies. (A) 3.4 Å cryo-EM density for class 2 C548-S complex (only the V_H_-V_L_ domains of C548 are shown). (B) Close-up view of quaternary epitope involving bridging interactions between adjacent RBDs. (C) CDR loops mapped onto adjacent RBD surfaces. (D) Epitope of C548 highlighted on adjacent RBDs. (E) C548 paratope mapped onto adjacent RBDs. (F) Interactions between RBD and C548 with a subset of interacting residues highlighted as sticks. Potential hydrogen bonds shown as dotted lines. (G) 3.4 Å cryo-EM density for class 3 C032-S complex (only the V_H_-V_L_ domains of C032 are shown). (H) Overlay of C032–RBD portion of the C032-S complex structure with an ACE2-RBD structure (from PDB 6VW1). (I) Epitope of C032 highlighted on the RBD surface. (J) C032 paratope mapped onto RBD surface. (K) Interactions between RBD and C032 CDRH1 and CDRH3 loops, with a subset of interacting residues highlighted as sticks. Potential hydrogen bonds shown as dotted lines. See also Figure S5, S7

To derive global properties of Fab-antigen interactions, we calculated shape complementarity (*Sc*) indices, which vary from 0 (not complementary) to 1 (a perfect fit) and are typically 0.64-0.68 for antibody-antigen interfaces (Lawrence and Colman, 1993). For antibody pairs for which we had determined both 1.3m and 6.2m structures, *Sc* values for 6.2m antibodies were modestly increased compared with their 1.3m counterparts: 0.56 versus 0.52 for C051 and C144 complexes with Spike, respectively (*Sc* values calculated for a Fab complexed with two adjacent RBDs), and 0.73 and 0.68 for C099 and C098 Fab complexes with RBD, respectively. Similarly, buried surface area (BSA) calculations did not reveal large increases in Fab-antigen interface areas upon antibody maturation: total BSAs for C051 and C144 interfaces were ∼2520 Å^2^ and ∼2350 Å^2^, respectively, and ∼2540 Å^2^ and ∼2590 Å^2^ for C099 and C098, respectively.

To understand the influence of individual mutations on potency and viral escape, we aligned RBD-bound Fab complexes from clonally-related 1.3m and 6.2m antibodies and inspected residue-level antibody-antigen interactions (Figure 6 and Figure S6). For the class 1 C098/C099 lineage, we compared 2.0 Å and 2.6 Å X-ray structures of the C098-RBD and C099-CR3022-RBD complexes, respectively (Figure S5H,I; Table S2). After superimposing the RBDs, the Fab V_H_-V_L_ domains adopted the same binding pose such that the CDR loops at the Fab-RBD interface were aligned equivalently (Figure 6A,B). Overall, the footprints of the epitope on the RBD and the paratope on the Fab were conserved (Figure S6A-C), consistent with the highly similar binding orientations of class 1 anti-RBD neutralizing antibodies (Figure S6D).

For C098 and C099, the majority of RBD contacts are mediated by CDR1 and CDR2 V gene-encoded regions (Figure S6A-C). Given that the C098 V_H_ and V_L_ gene segment sequences contained no somatic hypermutations (Figure S6A), our structures provided the opportunity to analyze the effects of affinity maturation on the increased potency of the 6.2m C099 antibody. Somatic mutations in C099 occurred in V gene-encoded CDR loops and FWRs, while the CDR3 loops remained unchanged from the germline (C098) antibody (Figure S6A,E). As previously noted for class 1 anti-RBD neutralizing antibodies (Hurlburt et al., 2020; Tan et al., 2021) somatic mutations in C099’s CDRH1 and CDRH2 appeared to drive improved binding and neutralizing characteristics. For example, the F27I_HC_ mutation found in C099 introduces a smaller hydrophobic residue that likely makes the CDRH1 loop more flexible, facilitating increased polar contacts and van der Waals interactions in this region (Figure 6C,D). Interestingly, in CDRH2, somatic mutations S53A_HC_ and Y58F_HC_ remove polar contacts with backbone carbonyl and side chain atoms at the RBD interface (Figure 6E,F). Yet, these mutations (particularly Y58F_HC_) increase binding affinity and neutralizing activity of class 1 anti-RBD antibodies (Tan et al., 2021), which can be partly explained by the introduction of stacking interactions with RBD residue T415 (Figure 6F). Thus, we conclude that a set of common somatic mutations found in C099 facilitates its improved neutralization potency.

For the class 2 C144/C051 lineage, we compared our previously-reported 3.2 Å cryo-EM structure of a C144 Fab-S complex (Barnes et al., 2020b) with the 3.5 Å C051-S structure reported here (Figure 6G). The C144 and C051 Fabs associate with the RBD through a similar binding mode to bridge adjacent RBDs on the surface of the S trimer (Figure 6H). As with C144, the C051 antibody heavy chain mediated the majority of RBD contacts (Figure S6F-H). Mutations at RBD positions L455, F456, E484, and Q493 conferred escape from C144, while only the E484K mutation conferred escape from C051 (Figure 1A). Viral escape at RBD positions L455 and Q493 is facilitated by an arginine substitution that would disrupt hydrogen-bonding networks at the C144-RBD interface (Figure 6I). Somatic mutations in the C051 CDRH1 (T28G_HC_ and S30R_HC_) introduce new polar contacts with backbone carbonyl and side chain residues at the RBD interface, while allowing additional flexibility in CDRH1 (Figure 6J), similar to observations for class 1 antibodies (Figure 6D). In addition, the CDRH3 E97D_HC_ somatic mutation in C051 introduces a smaller charged residue that may better accommodate an arginine side chain in this region (Figure 6J). Somatic hypermutations in CDRH1 for this lineage likely play an important role, as the clonally-related C054 antibody isolated at 6.2m is sensitive to the Q493R and L455R mutations (Gaebler et al., 2021).

### Structures of Spike trimer complexes with 1.3m class 2 and class 3 antibodies explain viral escape

To further understand escape patterns of RBD-targeting antibodies, we determined cryo-EM structures of Fab-S complexes for 1.3m class 2 (C548) and class 3 (C032) neutralizing antibodies (Figure 7 and Figure S5) and derived homology models of the 6.2m counterparts: C549 and C080, respectively (Figure S7). In both experimentally-determined structures, Fabs recognized either “up” or “down” RBD conformations (Figure 7).

The 3.4 Å cryo-EM structure of C548 Fabs bound to a closed S trimer (Figure 7A) revealed a quaternary epitope that spanned neighboring RBDs (Figure 7B,C). The antibody paratope involved five of six CDR loops, with the majority of contacts to RBD focused on residues involved in ACE2 recognition (Figure 7D and Figure S7). C548 is encoded by the *VH1-69* V_H_ gene segment, which encodes a hydrophobic sequence at the tip of CDRH2 that has been shown to facilitate broad neutralization against influenza, Hepatitis C, and HIV-1 (Chen et al., 2019). In C548, residues I53-F54_HC_ target a hydrophobic patch in the neighboring RBD core that resides near the base of the N343-glycan and comprises RBD residues W436, N440, and L441 (Figure 7D,E). These interactions are akin to those observed in the C144/C051 lineage, in which either Phe-Trp or Leu-Trp at the tip of CDRH3 is buried in a similar manner on the adjacent RBD (Figure S6G,H). These data demonstrate convergent evolution of mechanisms for anti-RBD antibodies to target this hydrophobic patch on the RBD surface, with the potential to lock RBDs into the “down” position.

Viral escape from C548 was mediated by substitutions at positions L455, E484, F490 and Q493, likely due to the disruption of polar contacts at the RBD interface and/or insertion of bulky sidechains into a sterically-restricted region (Figure 7F). However, unlike the C144/C051 lineage, C549 (the 6.2m mature counterpart) shows resilience to all C548 viral escape mutants, including partial resilience to the E484K substitution (Figure 1D). C549 exhibits accumulated somatic mutations (9 HC residues and 11 LC residues changed compared to germline) in both FWR and CDR loops (Figure S7A). Using the C548-S structure, we made a homology model of the C549-RBD interactions (Figure S7B). Light chain somatic mutations are predicted to explain the increased resistance: 30Y_LC_ provides additional stacking interactions with RBD residue F490, while 27D_LC_ and 92E_LC_ increase polar contacts with RBD backbone carbonyl and side chain atoms (Figure S7C). Thus, we predict that C549 partial resilience to the E484K mutation is not the result in changes in direct interactions with the E484 sidechain, but rather a series of adjacent residue changes to accommodate the E484K substitution.

To understand maturation in the C032/C080 class 3 antibody lineage, we determined a 3.3 Å cryo-EM structure of a C032-S trimer complex, revealing a Fab binding orientation that does not overlap with the ACE2 binding site (Figure 7G,H). C032 recognizes a glycopeptidic epitope focused on a short helical segment in the RBD core that spans RBD residues 437-442 near the N343-glycan (Figure 7I), with a paratope BSA (∼810 Å^2^) equally distributed among the CDRH1, CDRH2, CDRH3, and CDRL3 loops (Figure 7J and Figure S7A). At the tip of CDRH3, hydrophobic residues A97_HC_, V98_HC_, and W100_HC_ bury into a pocket shaped by RBD loops comprising residues 344-348 and 443-450 (Figure 7K), providing sequence-independent van der Waals interactions with the RBD backbone. Mutation of residues comprising this RBD pocket confer C032-resistance (Figure 3E). To predict how the affinity matured 6.2m antibody, C080, avoids viral escape, we made a homology model of the C080-RBD structure. The majority of somatic mutations in C080 are positioned distal to the modeled Fab-RBD interface (Figure S7D). Thus, it is likely that C080 somatic mutations influence CDR loop conformation and flexibility at the antigen interface, as has been observed for neutralizing antibodies against the HIV-1 Envelope (Klein et al., 2013). Interestingly, C080 also acquired activity against SARS-CoV (Figure 5F). Based on the homology model of C080-SARS-CoV RBD, CDRL3 mutations are predicted to facilitate recognition of the SARS-CoV RBD (Figure S7E).

## DISCUSSION

Herein we describe properties of SARS-CoV-2 neutralizing antibodies that change as a consequence accumulated somatic mutations in convalescent individuals (Gaebler et al., 2021; Sokal et al., 2021). Persistent somatic mutation is associated with continued availability of antigen (Victora and Nussenzweig, 2012). For example, during chronic HIV-1 infection, antibodies develop exceptionally large numbers of mutations compared to infections of limited duration (Klein et al., 2013; Scheid et al., 2009). In SARS-CoV-2 convalescent individuals, viral proteins and nucleic acids can persist in the gut for months, providing a source of antigen to fuel germinal centers (Gaebler et al., 2021). Whether current vaccination schemes will afford a sufficient antigen persistence to elicit continued antibody maturation remains to be determined.

While each antibody lineage had unique characteristics that were impacted by somatic mutations, general themes were evident. Typically, antibodies isolated at 6.2m had increased potency compared to their clonal relatives isolated at 1.3m. An exception to this was C144, a particularly potent antibody, isolated at 1.3m, (Robbiani et al., 2020). Structural analysis suggests that the high potency of C144 is related to its ability to lock the S trimer in a prefusion, closed state (Barnes et al., 2020b).

Whereas antibody producing plasma cells are selected based on their affinity for antigen, memory B-cells are heterogeneous and encode a far more diverse set of antibodies with varying levels of affinity for the immunogen (Viant et al., 2020). One of the consequences of accumulating a diverse group of closely related antibody-producing cells in the memory compartment is the ability to recognize and respond to closely related pathogens (Viant et al., 2020). Consistent with this observation, an important property that was recurrently evident in the antibody lineages described herein was a change in the mutations that were selected and conferred resistance to 6.2m antibodies as compared to 1.3m antibodies. Striking features of some of the 6.2m antibodies included restriction of the range of options for viral escape and the resilience of neutralization activity in the face of point mutations that conferred resistance to 1.3m antibodies. Indeed, the neutralization potency of certain matured antibodies, such as C549, C099 and C080, was maintained for all of the naturally circulating individual RBD substitutions tested, consistent with observation of antibody antigen structures or models. In some cases, rVSV/SARS-CoV-2 selection experiments indicated that somatic mutations elevated the genetic barrier to antibody resistance, imposing a requirement for two substitutions for high level antibody resistance.

The naturally circulating RBD triple mutant K417N/E484K/N501Y did not generally confer resistance to antibodies that were not already affected by the E484K mutation. This finding suggests that separate antibodies may be generally responsible for the application of selection pressure at K417, E484 and N501. Nevertheless, the E484K mutation undermined the activity of several class 2 antibodies. While a number of naturally circulating substitutions at E484 conferred resistance to some class 2 antibodies (e.g., C144, C055, C548), naturally occurring variants of concern typically encode E484K (West et al., 2021), consistent with our finding that only the E484K substitution conferred more pervasive class 2 antibody resistance, including to some matured antibodies (e.g., C051, C052).

In two cases, antibody maturation enabled neutralization of heterologous sarbecoviruses, suggesting that development of pan-sarbecovirus vaccines may be possible (Cohen et al., 2021b). Indeed, the greater neutralization potency, resilience to viral mutation, and breadth of SARS-CoV-2 RBD-specific antibodies that have undergone greater degrees of somatic mutation suggests that immunization schemes that elicit higher levels of antibody mutation and diversification are desirable. Indeed, antibody maturation may be especially important as SARS-CoV-2 diversifies and adapts to the range of human antibodies in vaccinated and previously infected individuals. Moreover, a diverse set of broadly neutralizing SARS-CoV-2 spike-elicited antibodies that exhibit some activity against divergent sarbecoviruses may mitigate the threat posed by this group of pandemic-threat agents.

## Acknowledgements

We thank all study participants and clinical staff. We thank members of the Bjorkman, Nussenzweig, and Bieniasz laboratories for helpful discussions, Dr. Jost Vielmetter, Pauline Hoffman, and the Protein Expression Center in the Beckman Institute at Caltech for expression assistance. Electron microscopy was performed in the Caltech Beckman Institute Resource Center for Transmission Electron Microscopy with assistance from Dr. Songye Chen. We thank the Gordon and Betty Moore and Beckman Foundations for gifts to Caltech to support the Molecular Observatory (Dr. Jens Kaiser, director), and Drs. Silvia Russi, Aina Cohen, and Clyde Smith and the beamline staff at SSRL for data collection assistance. Use of the Stanford Synchrotron Radiation Lightsource, SLAC National Accelerator Laboratory, is supported by the U.S. Department of Energy, Office of Science, Office of Basic Energy Sciences under Contract No. DE-AC02-c76SF00515. The SSRL Structural Molecular Biology Program is supported by the DOE Office of Biological and Environmental Research, and by the National Institutes of Health, National Institute of General Medical Sciences (P41GM103393). This work was supported by NIH grants R37-AI64003 (to P.D.B.), R01AI78788 (to T.H.), P01-AI138938-S1 (P.J.B. and M.C.N.), K99 AI153465 (to A.I.F.), 2U19AI111825 (to M.C.N.). This work was also supported by a George Mason University Fast Grant (P.J.B.), a grant from the NSF GRFP DGE-1745301 (to A.T.D.), and by the Caltech Merkin Institute for Translational Research (P.J.B.). C.O.B was supported by the Hanna Gray Fellowship Program from the Howard Hughes Medical Institute and the Postdoctoral Enrichment Program from the Burroughs Wellcome Fund. F.M. was supported by the Bulgari Women & Science Fellowship in COVID-19 Research. C.G. was supported by the Robert S. Wennett Post-Doctoral Fellowship, in part by the National Center for Advancing Translational Sciences (National Institutes of Health Clinical and Translational Science Award program, grant UL1 TR001866), and by the Shapiro-Silverberg Fund for the Advancement of Translational Research. M.C.N. and P.D.B. are Howard Hughes Medical Institute Investigators. The contents of this publication are solely the responsibility of the authors and do not necessarily represent the official views of NIGMS, NIAID or NIH.

## Author contributions

F.M., Y.W., C.O.B., F.S., M.C.N, P.J.B., T.H., and P.D.B., conceived the study and analyzed data. F.S. and J.D.S. generated spike plasmids. F.M., D.S.B., J.C.L and J.D.S. performed neutralization assays with natural SARS-CoV-2 variant and sarbecovirus HIV-1 pseudotypes. Y.W., F.S., and M.R. performed the selection and characterization of escape mutants using VSV/SARS-CoV-2 and HIV-1-pseudotypes. C.O.B. and K.H.T. performed protein purification and complex assembly. A.I.F. and C.O.B. performed crystallographic studies and analyzed structures. C.O.B., A.T.D., and A.I.F. performed cryo-EM studies and analyzed structures. S.H. and C.A.S. generated and performed analyses of homology models. Z.W., S.F., A.C., T.Y.O., M.C., K.G.M., V.R. and A.G. isolated and characterized monoclonal antibodies. C.G. and M.C. recruited participants and executed clinical protocols. F.M., Y.W., C.O.B., F.S., M.C.N, P.J.B., T.H. and P.D.B. wrote the paper with contributions from other authors.

## LEAD CONTACT AND MATERIALS AVAILABILITY

Requests for further information and or reagents may be addressed to the Lead Contact Paul D. Bieniasz.

## METHODS DETAILS

### SARS-CoV-2 pseudotyped reporter virus

We have previously described a panel of plasmids expressing RBD-mutant SAR-CoV-2 spike proteins in the context of pSARS-CoV-2-SΔ19 (Weisblum et al., 2020). Additional substitutions to expend the panel were introduced using synthetic gene fragments (IDT) or overlap extension PCR mediated mutagenesis and Gibson assembly. E484K was originally excluded from our panel because HIV-1-based pseudotypes generated with the E484K substitution in our standard assay were poorly infectious. However, when the E484K substitution was incorporated into a spike protein that also includes that the R683G substitution, which disrupts the furin cleavage site, pseudotyped particle infectivity was preserved. The R683G substitution itself increased pseudovirus sensitivity to some antibodies, including C055, C099, C549 and C512, and antibodies from the C144 and C032 groups. Thus, the E484K, L455R+E484K and KEN (K417N+E484K+N501Y) mutants were used in the context of a pSARS-CoV-2-S Δ19 variant with an inactivated furin cleavage site (R683G). The potencies with which the antibodies neutralized members of the mutant pseudotype panel were compared with potencies against a “wildtype” SARS-CoV-2 (NC_045512) spike sequence, carrying R683G where appropriate. The SARS-CoV-2 pseudotyped HIV-1 particles were generated as previously described (Schmidt et al., 2020). Specifically, virus stocks were harvested 48 hours after transfection of 293T cells with pNL4-3ΔEnv-nanoluc and pSARS-CoV-2 SΔ19 and filtered and stored at −80°C.

### SARS-CoV-2 pseudotype neutralization assays

Monoclonal antibodies were four-fold serially diluted and then incubated with SARS-CoV-2 pseudotyped HIV-1 reporter virus for 1 h at 37 °C. The antibody/pseudotyped virus mixture was then added to HT1080/ACE2.cl14 cells. After 48 h, cells were washed with PBS, lysed with Luciferase Cell Culture Lysis reagent (Promega) and Nanoluc Luciferase activity in lysates was measured using the Nano-Glo Luciferase Assay System (Promega) and a Glomax Navigator luminometer (Promega). The relative luminescence units were normalized to those derived from cells infected with SARS-CoV-2 pseudotyped virus in the absence of monoclonal antibodies. The half-maximal inhibitory concentrations for monoclonal antibodies (IC_50_) were determined using four-parameter nonlinear regression (least squares regression method without weighting) (GraphPad Prism).

### Selection of antibody resistant rVSV/SARS-CoV-2 variants

To select monoclonal antibody-resistant S variants, rVSV/SARS-CoV-2/GFP_1D7_ and rVSV/SARS-CoV-2/GFP_2E1_ were passaged to generate diversity, and populations containing 10^6^ PFU were incubated with monoclonal antibodies (0.5µg/ml to 10µg/ml) for 1h at 37°C before inoculation of 2×10^5^ 293T/ACE2cl.22 cells in 6-well plates. The following day the medium was replaced with fresh medium containing the same concentrations of antibody. Supernatant from the wells containing the highest concentration of monoclonal antibodies that showed evidence of rVSV/SARS-CoV-2/GFP replication (large numbers of GFP positive cells or GFP positive foci) was harvested 24h later. Where necessary, aliquots (100 µl) of the cleared supernatant from the first passage (p1) were incubated with the same concentration of monoclonal antibody and then used to infect 2×10^5^ 293T/ACE2cl.22 cells in 6-well plates, as before (p2). We repeated this process until escape reduced neutralization potency for the antibody was evident, as indicated by increasing numbers of GFP positive cells.

To isolate individual mutant viruses by limiting dilution, the selected rVSV/SARS-CoV-2/GFP_1D7_ and rVSV/SARS-CoV-2/GFP_2E1_ populations were serially diluted in the absence of monoclonal antibodies and aliquots of each dilution added to individual wells of 96-well plates containing 1×10^4^ 293T/ACE2cl.22 cells. Individual viral variants were identified by observing single GFP-positive plaques at limiting dilutions. The plaque-purified viruses were expanded, RNA extracted and S sequences determined, and sensitivity to the selecting monoclonal antibody measured.

### rVSV/SARS-CoV-2 Neutralization assays

Monoclonal antibodies were five-fold serially diluted and then incubated with rVSV/SARS-CoV-2/GFP_1D7_ and rVSV/SARS-CoV-2/GFP_2E1_ or plaque purified selected variants for 1 h at 37 °C. The antibody/recombinant virus mixture was then added to 293T/ACE2.cl22 cells. After 16h, cells were harvested, and infected cells were quantified by flow cytometry. The percentage of infected cells was normalized to that derived from cells infected with rVSV/SARS-CoV-2 in the absence of monoclonal antibodies. The half-maximal inhibitory concentrations for monoclonal antibodies (IC_50_) were determined using four-parameter nonlinear regression (least squares regression method without weighting) (GraphPad Prism).

### Sequence analyses

To identify putative antibody resistance mutations, RNA was isolated from aliquots of supernatant containing selected viral populations or individual plaque purified variants using NucleoSpin 96 Virus Core Kit (Macherey-Nagel). The purified RNA was subjected to reverse transcription using random hexamer primers and SuperScript VILO cDNA Synthesis Kit (Thermo Fisher Scientific). The cDNA was amplified using KOD Xtreme Hot Start DNA 396 Polymerase (Millipore Sigma) flanking the S encoding sequences. Alternatively, a fragment including the entire S-encoding sequence was amplified using primers targeting VSV-M and VSV-L. The PCR products were gel-purified and sequenced either using Sanger-sequencing or Illumina sequencing as previously described (Gaebler et al., 2019). For illumina sequencing, 1µl of diluted DNA was used with 0.25 µl Nextera TDE1 Tagment DNA enzyme (catalog no. 15027865), and 1.25 µl TD Tagment DNA buffer (catalog no. 15027866; Illumina). Then, the DNA was ligated to i5/i7 barcoded primers using the Illumina Nextera XT Index Kit v2 and KAPA HiFi HotStart ReadyMix (2X; KAPA Biosystems). Next the DNA was purified using AmPure Beads XP (Agencourt), pooled, sequenced (paired end) using Illumina MiSeq Nano 300 V2 cycle kits (Illumina) at a concentration of 12pM. For analysis of the Illumina sequencing data, adapter sequences were removed from the raw reads and low-quality reads (Phred quality score <20) using BBDuk. Filtered reads were mapped to the codon-optimized SARS-CoV-2 S sequence in rVSV/SARS-CoV-2/GFP and mutations were annotated using using Geneious Prime (Version 2020.1.2), using a P-value cutoff of 10^-6^. RBD-specific variant frequencies, P-values, and read depth were compiled using Python running pandas (1.0.5), numpy (1.18.5), and matplotlib (3.2.2).

### Protein expression and purification

Expression and purification of SARS-CoV-2 6P stabilized S trimers (Hsieh et al., 2020) and SARS-CoV-2 RBD were conducted as previously described (Cohen et al., 2021a). We purified proteins from supernatants of transiently-transfected Expi293F cells (Gibco) by Ni^2+^-NTA affinity and size exclusion chromatography (SEC). Peak fractions from SEC were identified by SDS-PAGE, pooled, and stored at 4°C. Fabs were generated by papain digestion from purified IgGs using crystallized papain (Sigma-Aldrich) in 50 mM sodium phosphate, 2 mM EDTA, 10 mM L-cysteine, pH 7.4 for 30-60 min at 37°C at a 1:100 enzyme:IgG ratio. To remove Fc fragments and undigested IgGs, digested products were applied to a 1-mL HiTrap MabSelect SuRe column (GE Healthcare Life Sciences) and the flow-through containing cleaved Fabs was collected. Fabs were further purified by SEC using a Superdex 200 Increase 10/300 column (GE Healthcare Life Sciences) in TBS before concentrating and storing at 4°C.

### Cryo-EM structure determinations

We incubated purified Fab and S 6P trimer at a 1.1:1 molar ratio per protomer on ice for 30 minutes prior to deposition on a freshly glow-discharged 300 mesh, 1.2/1.3 UltrAuFoil grid or 1.2/1.3 QuantiFoil Cu grid. Fluorinated octyl-maltoside was added to the Fab-S complex to a final detergent concentration of 0.02% w/v, resulting in a final complex concentration of 3 mg/ml, immediately before 3 µl of complex was applied to the grid. Samples were then vitrified in 100% liquid ethane using a Mark IV Vitrobot after blotting for 3 s with Whatman No. 1 filter paper at 22°C, 100% humidity.

We followed previously-described cryo-EM data collection and processing protocols for Fab-S complexes (Barnes et al., 2020a). Briefly, for all Fab-S complexes, we collected micrographs on a Talos Arctica transmission electron microscope (Thermo Fisher) operating at 200 kV using SerialEM automated data collection software (Mastronarde, 2005). Movies were recorded using a 3×3 beam image shift pattern with a K3 camera (Gatan). Data collection parameters are reported in **Table S3**. Cryo-EM movies were patch motion corrected for beam-induced motion including dose-weighting within cryoSPARC v2.15 (Punjani et al., 2017) after binning super resolution movies for all data sets. Non-dose-weighted images were used to estimate CTF parameters using a cryoSPARC implementation of the Patch CTF job, and all datasets were processed similarly. Briefly, after picking an initial set of particles based on templates from 2D classification of blob picked particles on a small sub-set of images, this set was pared down through several rounds of 3D classification. Ab initio cryoSPARC jobs on a small good subset of these particles revealed distinct states and junk particles. A full set of particles was heterogeneously refined against distinct conformational states and a junk class acting as a trap for bad particles. Particles from each class were separately refined using non-uniform refinement using C1 (C032-S) or C3 symmetry (C051-S and C548-S). Particles from distinct states were re-extracted without binning and were separately refined in rounds of 3D classification. Particles were further subdivided into groups based on beam-tilt and refined separately for CTF parameters and aberration correction. For the C032-S and C051-S data sets, a soft mask (3-pixel extension, 6-pixel soft edge) was generated for the spike S1 subunit and Fab variable domains to improve local resolutions at the Fab-RBD interface. Overall reported resolutions are based on gold standard FSC calculations (Scheres and Chen, 2012).

### Cryo-EM Structure Modeling and Refinement

Initial complex coordinates were generated by docking individual chains from reference structures into cryo-EM densities using UCSF Chimera (Goddard et al., 2018). Models were refined into cryo-EM densities using rigid body and real space refinement with morphing in Phenix (Terwilliger et al., 2018). Models with updated sequences were built manually in Coot (Emsley et al., 2010) and then refined using iterative rounds of real-space refinement in Phenix and Coot. *N*-Glycans were modeled at potential *N*-linked glycosylation sites (PNGSs) in Coot using ‘blurred’ maps processed with B-factors generated in cryoSPARC v2.15. We validated model coordinates using MolProbity (Chen et al., 2010) (**Table S3**).

### X-ray structures

To assemble C098-RBD or C099-CR3022-RBD complexes for crystallization, a 3:1 Fab:RBD molar ratio was incubated at RT for 1 h and complexes purified using size exclusion chromatography on a Superdex200 10/300 column (Cytiva) in 1x TBS. Crystallization trials for individual Fabs (C032, C080, C098, and C099), C098-RBD, and C099-CR3022-RBD complexes were carried out at room temperature using the sitting drop vapor diffusion method by mixing equal volumes of the Fab or Fab-RBD complex and reservoir using a TTP LabTech Mosquito robot and commercially-available screens (Hampton Research). C032 Fab crystals were grown using 0.2 µL of protein complex in TBS and 0.2 µL of mother liquor (0.1 M HEPES pH 7.7, 58% 2-Methyl-2,4-pentanediol) and cryoprotected in mother liquor. C080 Fab crystals were grown using 0.2 µL of protein complex in TBS and 0.2 µL of mother liquor (10% 2-Propanol, 0.1 M BICINE pH 8.5, 30% PEG 1,500) and cryoprotected using Fomblin® oil. C098 Fab crystals were grown using 0.2 µL of protein complex in TBS and 0.2 µL of mother liquor (2.0 M ammonium sulfate, citric acid pH 3.5) and cryoprotected in mother liquor supplemented with 15% (v/v) glycerol. C099 Fab crystals were grown using 0.2 µL of protein complex in TBS and 0.2 µL of mother liquor (1.9 M ammonium sulfate, citric acid pH 3.8) and cryoprotected using Al’s oil. C098-RBD crystals were grown using 0.2 µL of protein complex in TBS and 0.2 µL of mother liquor (0.05 M citric acid, 0.05M BIS-TRIS propane pH 5.0, 14% PEG 3,350) and cryoprotected in mother liquor supplemented with 30% (v/v) glycerol. C099-CR3022-RBD crystals were grown using 0.2 µL of protein complex in TBS and 0.2 µL of mother liquor (0.1M Sodium cacodylate, 40% 2-Methyl-2,4-pentanediol (MPD), and 5% PEG8000) and cryoprotected in mother liquor supplemented with 10% (v/v) glycerol.

X-ray diffraction data were collected for individual Fabs or Fab-RBD complexes at the Stanford Synchrotron Radiation Lightsource (SSRL) beamline 12-2 on a Pilatus 6M pixel detector (Dectris). Data from single crystals were indexed and integrated in XDS (Kabsch, 2010) or iMosflm {Battye, 2011 #796} and merged using AIMLESS in *CCP4* (Winn et al., 2011) (Table S2). Fab structures were solved by molecular replacement using a model of CC12.3 Fab (PDB6XC4) or HEPC46 Fab (PDB 6MEG). The C098-RBD complex structure was solved by molecular replacement using the C098 Fab (this paper) and RBD (PDB 7BZ5) structures as search models. The C099-CR3022-RBD complex structure was solved by molecular replacement using the C099 Fab (this paper), CR3022 Fabb (PDB 6W41) and RBD (PDB 7BZ5) structures as search models. Heavy chain and light chain CDR loops for the search model Fab were trimmed to make the search models. The structures were refined using an initial round of rigid body and individual B refinement in Phenix (Adams et al., 2010) followed by cycles of manual building in Coot (Emsley et al., 2010) and real space refinement in Phenix with TLS (**Table S2**).

### Homology modeling of Fab-RBD structures

Homology models of two of the RBD complexes were made for the Fabs of C549 (class 2) and C080 (class 3). Both complexes were modelled based on the cryo-EM structures of the related 1.3 month Fab-S complexes; i.e., the C548-S and C032-S complexes. The Fab of C548 to C549 involved 27 amino acid changes, whereas there were 16 changes for C032 to C080. Side chains of residues that were disordered in the density of the experimental structures were also modelled to their correct sequence. Homology models were generated by MODELLER (version 9.23) (Sali and Blundell, 1993) and further optimized by Protein Preparation Wizard in Maestro Schrodinger (Sastry et al., 2013) including optimization of hydrogens. The system was fully solvated with SPC water and counter ions. Energy minimization using Brownian Dynamics in Desmond from Schrodinger (version 2020.1) (Bowers et al., 2006) which involved gradually reducing restraints in 100ps steps from the full protein, to the backbone and finally without restraints to avoid any steric clashes.

### Structural Analyses

CDR definitions and Kabat numbering for antibody residues were based on IMGT definitions (Lefranc et al., 2015). Figures of structures were made with UCSF ChimeraX. Local resolution maps were calculated using cryoSPARC v 2.15. Areas buried in Fab-RBD interfaces (BSAs) were calculated using PDBePISA (Krissinel and Henrick, 2007) and a 1.4 Å probe. *Sc* analyses were conducted using Rosetta version 2020.08 (Leaver-Fay et al., 2011). Potential hydrogen bonds were assigned as interactions with A-D-H angle > 90° and between atoms that were <4.0Å. Potential van der Waals interactions were assigned as interactions that were <4.0Å. Hydrogen bond and van der Waals interaction assignments are tentative in the cryo-EM structures due to resolution limitations.

### DATA AND SOFTWARE AVAILABILITY

Coordinates and maps associated with data reported in this manuscript will be deposited in the Electron Microscopy Data Bank (EMDB: https://www.ebi.ac.uk/pdbe/emdb/) and Protein Data Bank (PDB: www.rcsb.org) with accession numbers …

## SUPPLEMENTAL INFORMATION

### Supplemental Figures S1-S7

**Figure S1.**
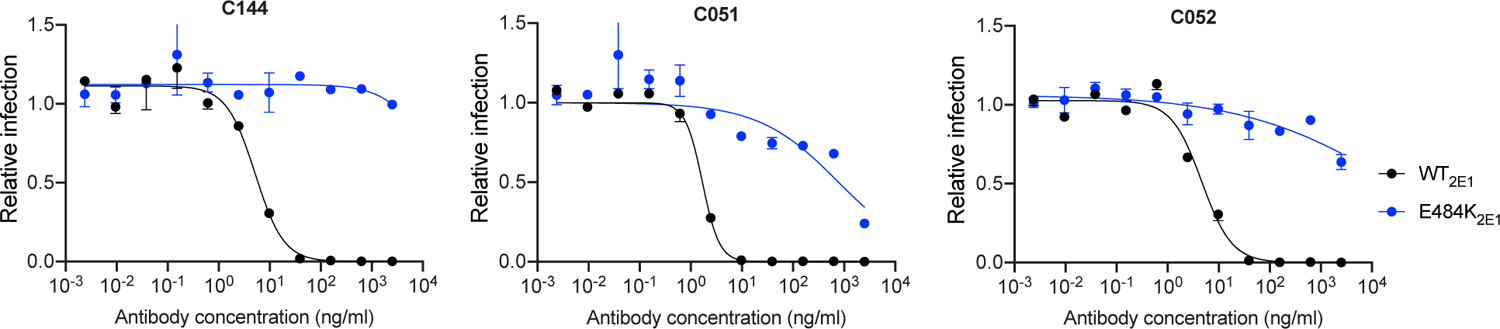
Effect of E484K substitution on C144/C051/C052 lineage neutralizing potency, related to Figure 1. Neutralization of rVSV/SARS-CoV-2 2E1 or a plaque purified E484K mutant thereof, in 293T/ACE2cl.22 cells. Infected (%GFP+) cells relative to no antibody controls, mean and range of two independent experiments is plotted.

**Figure S2.**
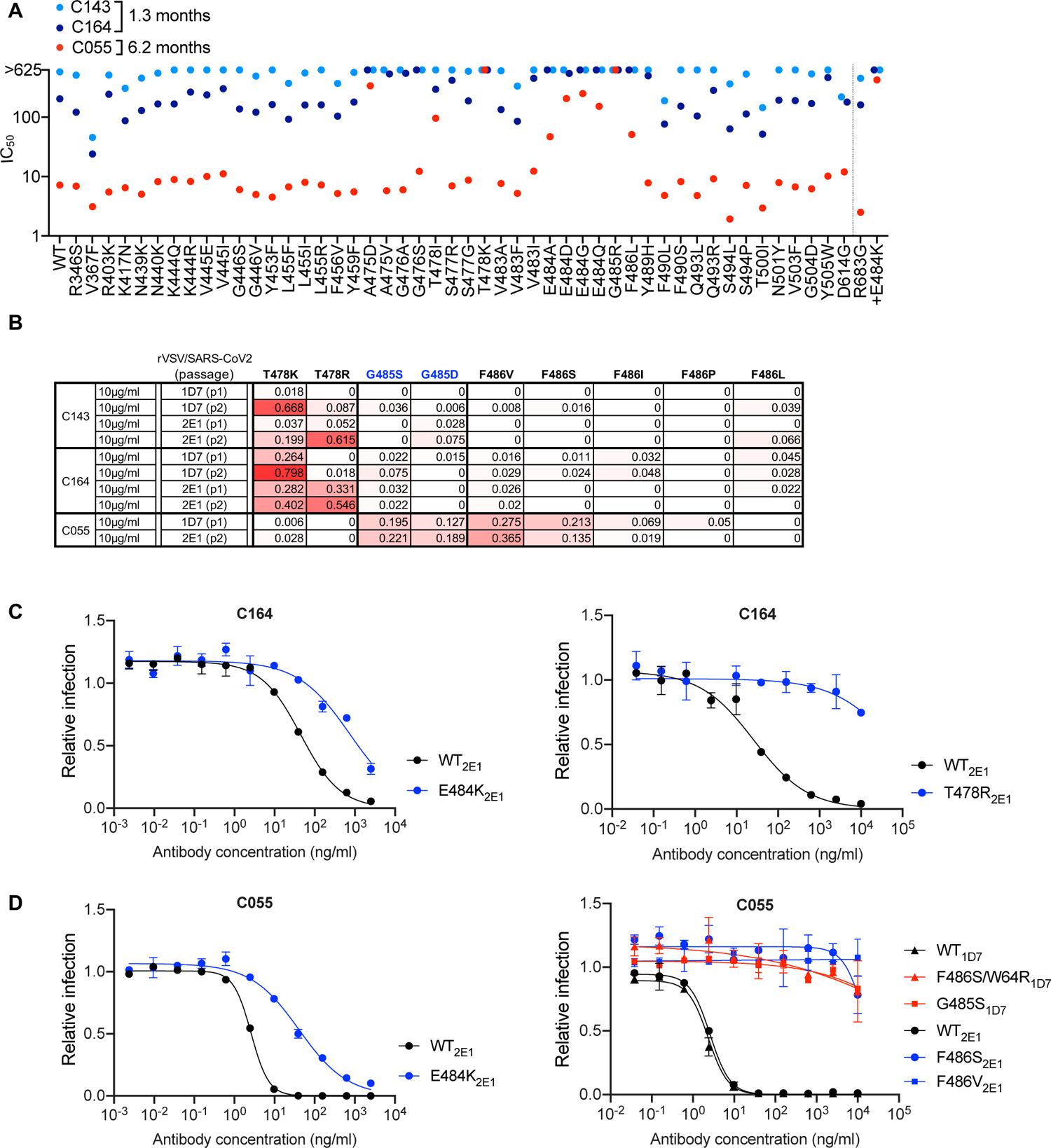
Effects of somatic mutation on potency and viral escape in the C143/C164/C055 class 2 antibody lineage, related to Figure 1. (A) Neutralization potency (IC_50_) of C143, C164 and C055 measured using HIV-1-based SARS-CoV-2 variant pseudotypes and HT1080/ACE2cl.14 cells. The E484K substitution was constructed in an R683G (furin cleavage site mutant) background to increase infectivity. Mean of two independent experiments. (B) Decimal fraction (color gradient; white = 0, red = 1) of Illumina sequence reads encoding the indicated RBD substitutions following rVSV/SARS-CoV-2 replication (1D7 and 2E1 virus isolates) in the presence of the indicated amounts of antibodies for the indicated number of passages. (C) C164 neutralization of rVSV/SARS-CoV-2 1D7, 2E1 or plaque purified mutants thereof, isolated following antibody selection, in 293T/ACE2cl.22 cells. Infected (%GFP+) cells relative to no antibody controls, mean and range of two independent experiments is plotted. (D) As in C for antibody C055.

**Figure S3.**
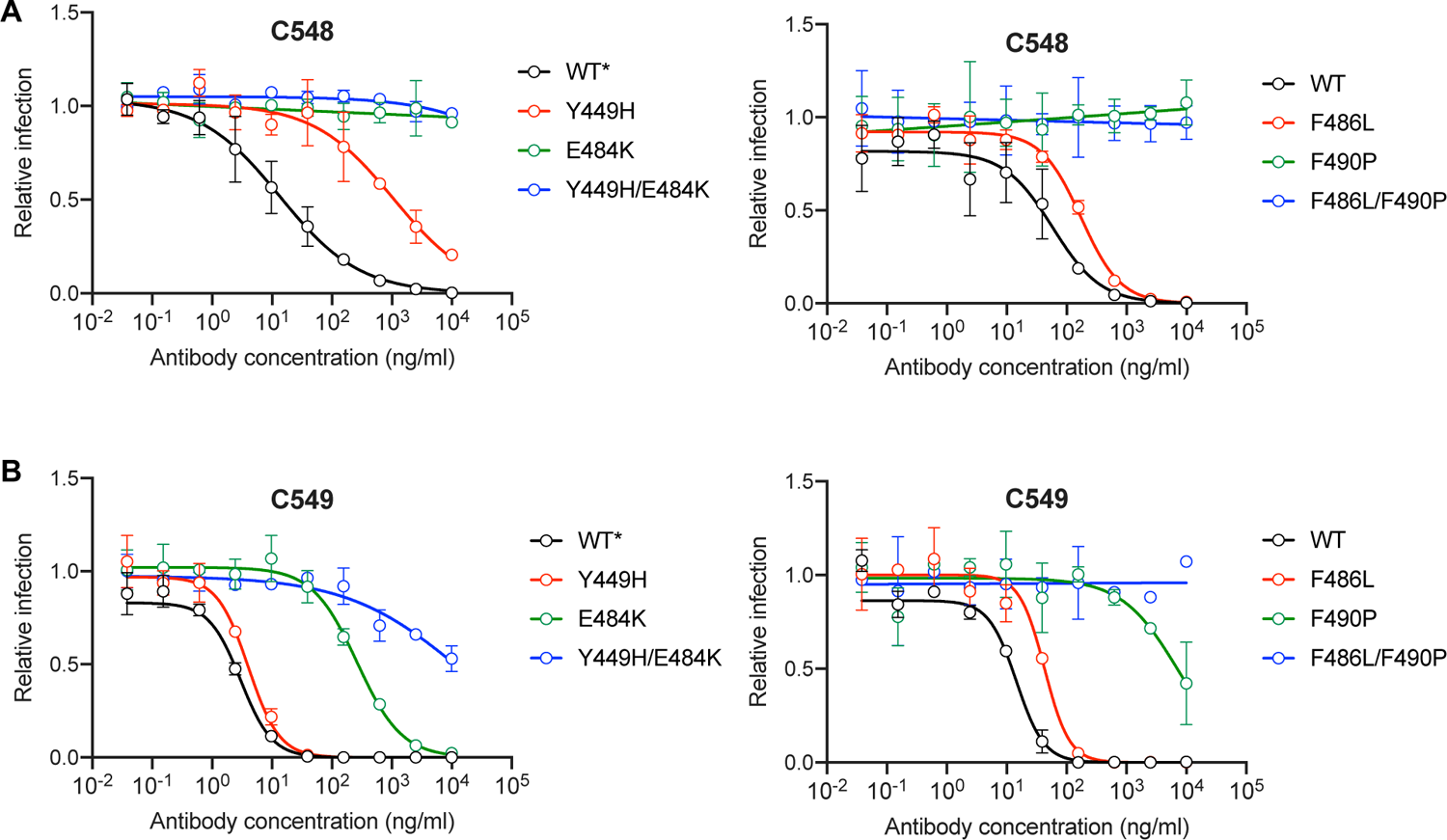
Effect of viral substitutions on neutralization by C548 and C549, related to Figure 1. (A) C548 neutralization of HIV-1-based SARS-CoV-2 pseudotypes harboring the indicated substitutions that were identified by selection experiments. Infection (NanoLuc luciferase activity) is normalized to that obtained in the absence of antibody, mean and range of two independent experiments is plotted. (B) As in A for antibody C549.

**Figure S4.**
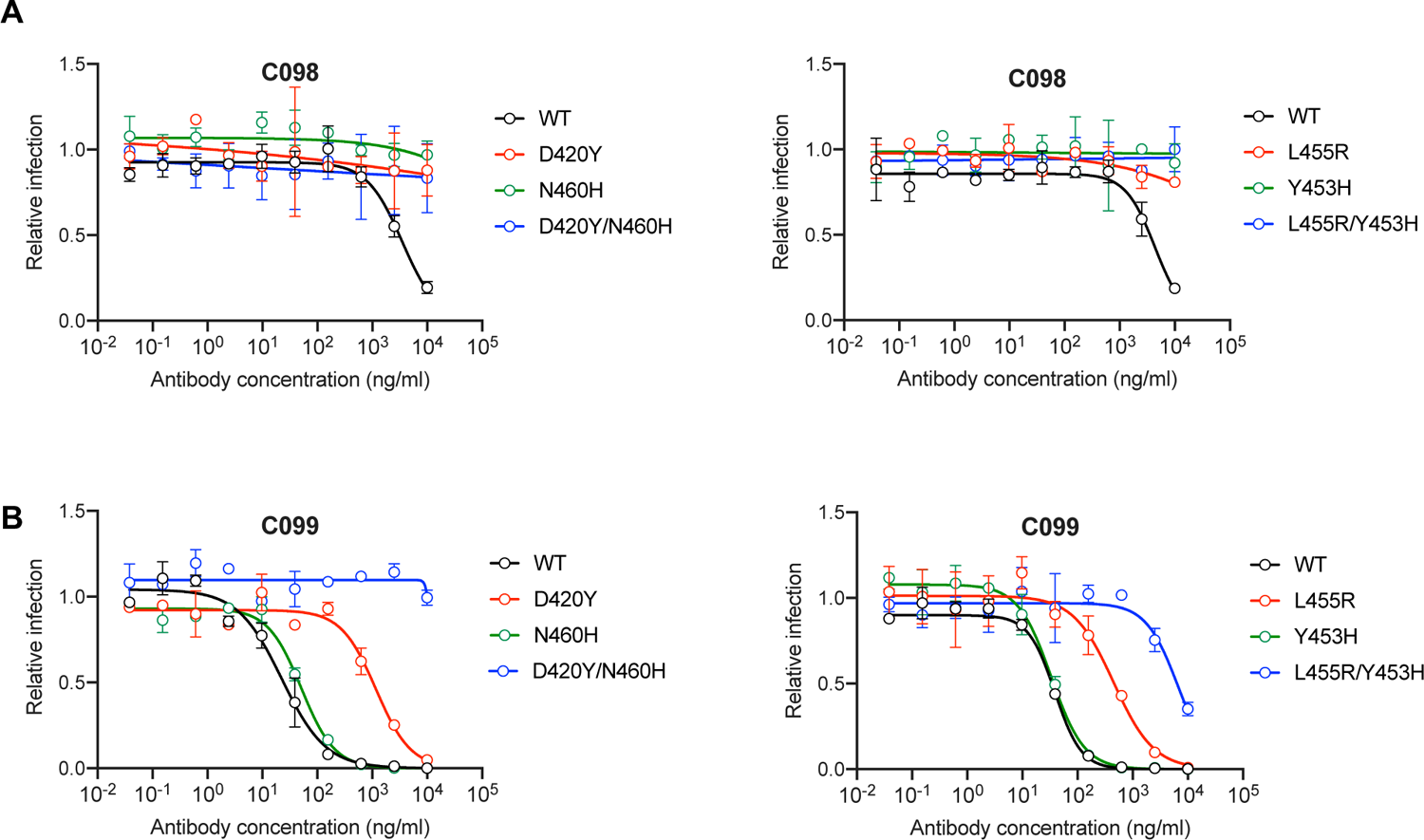
Effect of viral substitutions on neutralization by C098 and C099, related to Figure 2. (A) C098 neutralization of HIV-1-based SARS-CoV-2 pseudotypes harboring the indicated substitutions that were identified by selection experiments. Infection (NanoLuc luciferase activity) is normalized to that obtained in the absence of antibody, mean and range of two independent experiments is plotted. (B) As in A for antibody C099.

**Figure S5.**
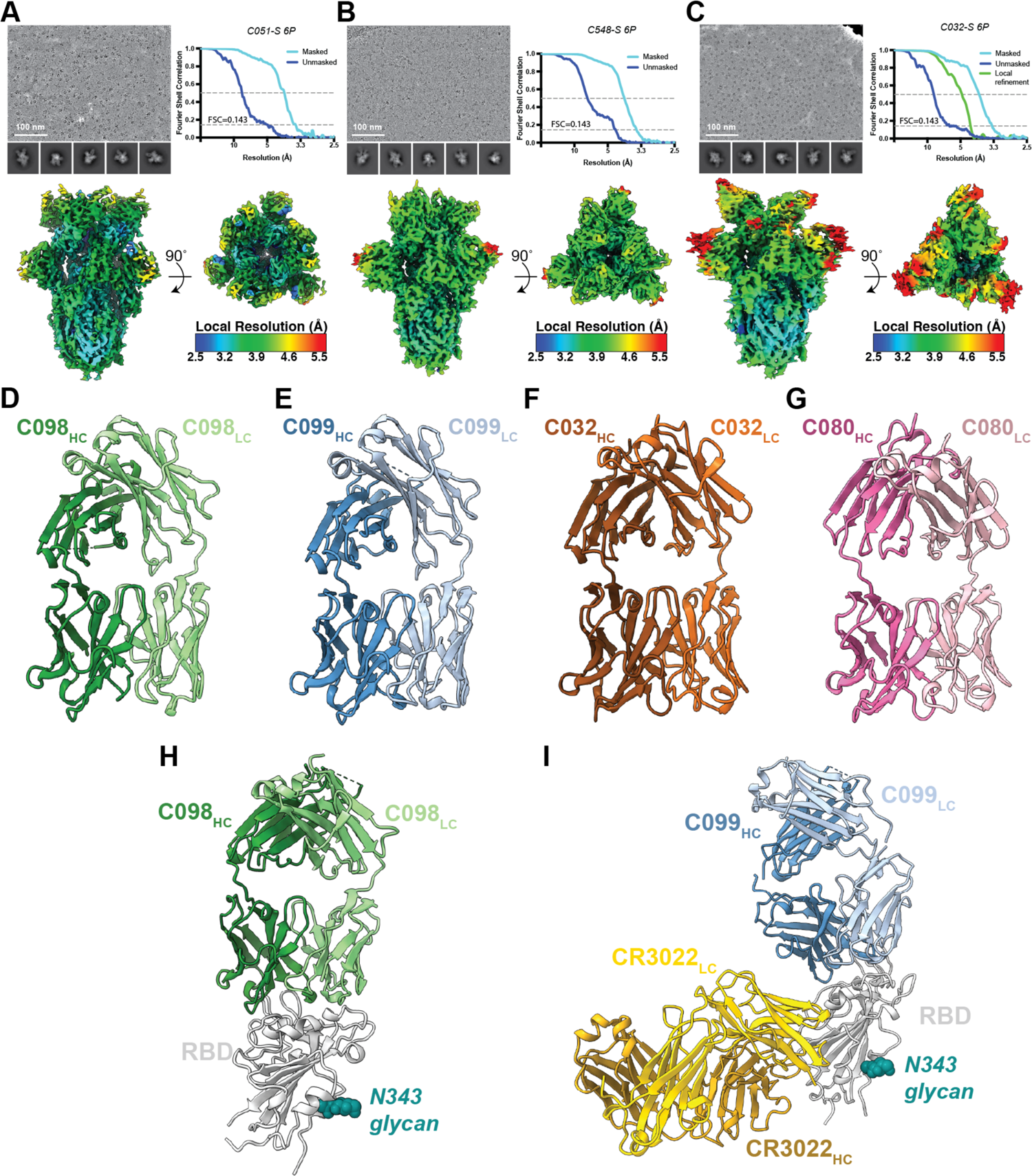
Cryo-EM data processing and X-ray structures, related to Figures 6,7. (A-C) Representative micrograph, 2D class averages, FSC plots calculated using the gold-standard FSC criteria, and local resolution maps rendered in cryoSparc v2.15 for the cryo-EM structures of (A) C051-S, (B) C032-S, and (C) C548-S complexes. (D-G) Cartoon representations of crystal structures of (D) C098, (E) C099, (F) C032, and (G) C080 Fabs. (H) Cartoon representation of C098 Fab – SARS-CoV-2 RBD crystal structure. (I) Cartoon representation of C099-CR3022 – SARS-CoV-2 RBD crystal structure.

**Figure S6.**
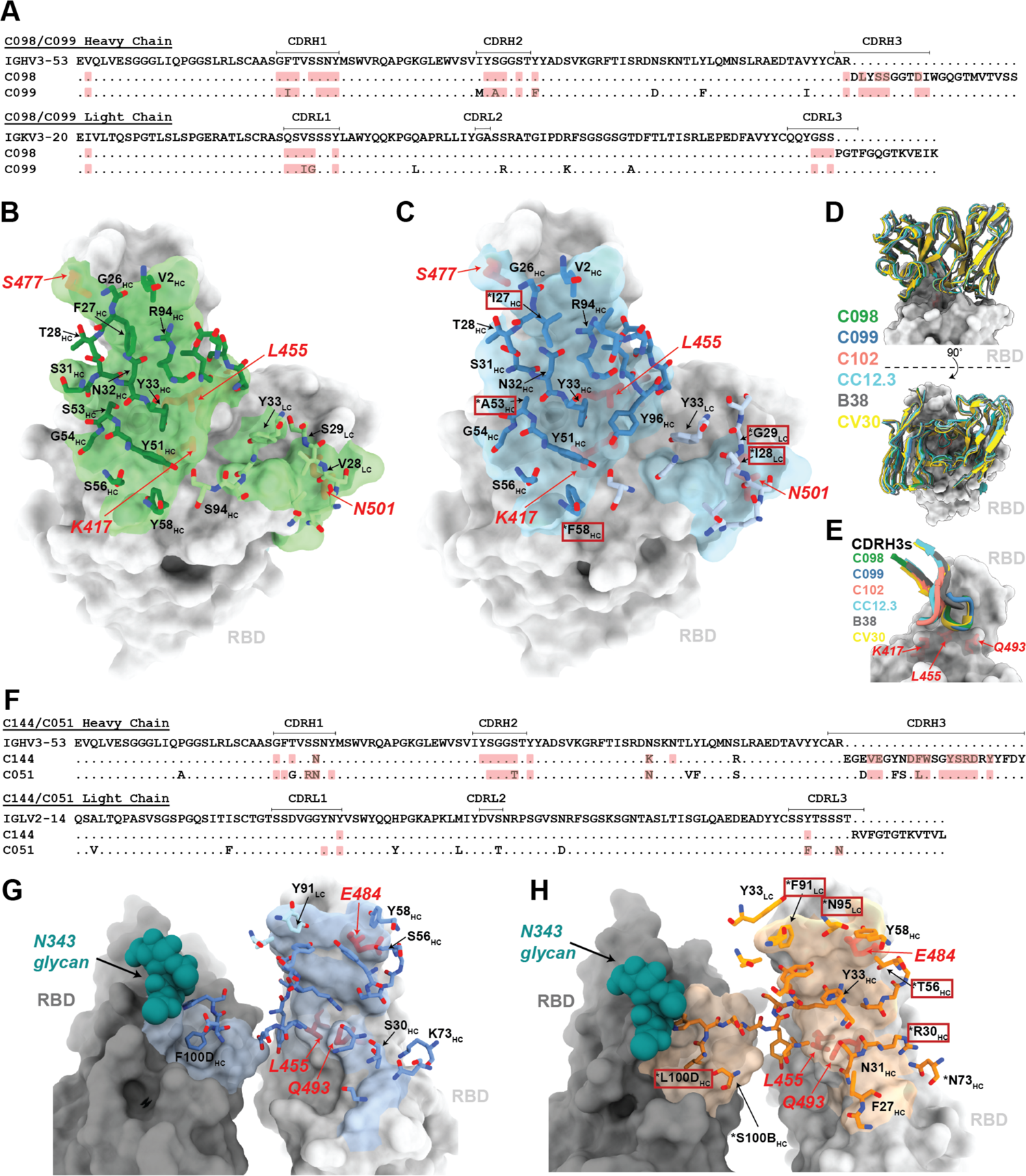
Class 1 and 2 antibody sequence alignments and interactions with RBD, related to Figure 6. (A) Sequence alignment between heavy and light chains of C098 and C099 relative to inferred germline sequences. Paratope residues highlighted in red. (B) C098 epitope (light green surface on RBD with paratope sidechains from C098 highlighted as sticks). (C) C099 epitope (light cyan surface on RBD with paratope sidechains from C099 highlighted as sticks). Somatic hypermutations found in C099 are highlighted with a red box. (D) Overlay of V_H_-V_L_ domains of class 1 Fabs bound to RBD (C098, green – this study; C099, blue – this study; C102, salmon – PDB 7K8M; CC12.3, cyan – PDB 6XC4; B38, gray – PDB 7BZ5; CV30, yellow – PDB 6XE1). (E) Overlay of CDRH3 loops of class 1 Fabs described in panel D at the RBD interface. (F) Sequence alignment between heavy and light chains of C144 and C051 relative to inferred germline sequences. Paratope residues highlighted in red. (G) C144 epitope (light blue surface on RBD with paratope sidechains from C144 highlighted as sticks). (H) C051 epitope (light orange surface on RBD with paratope sidechains from C051 highlighted as sticks).

**Figure S7.**
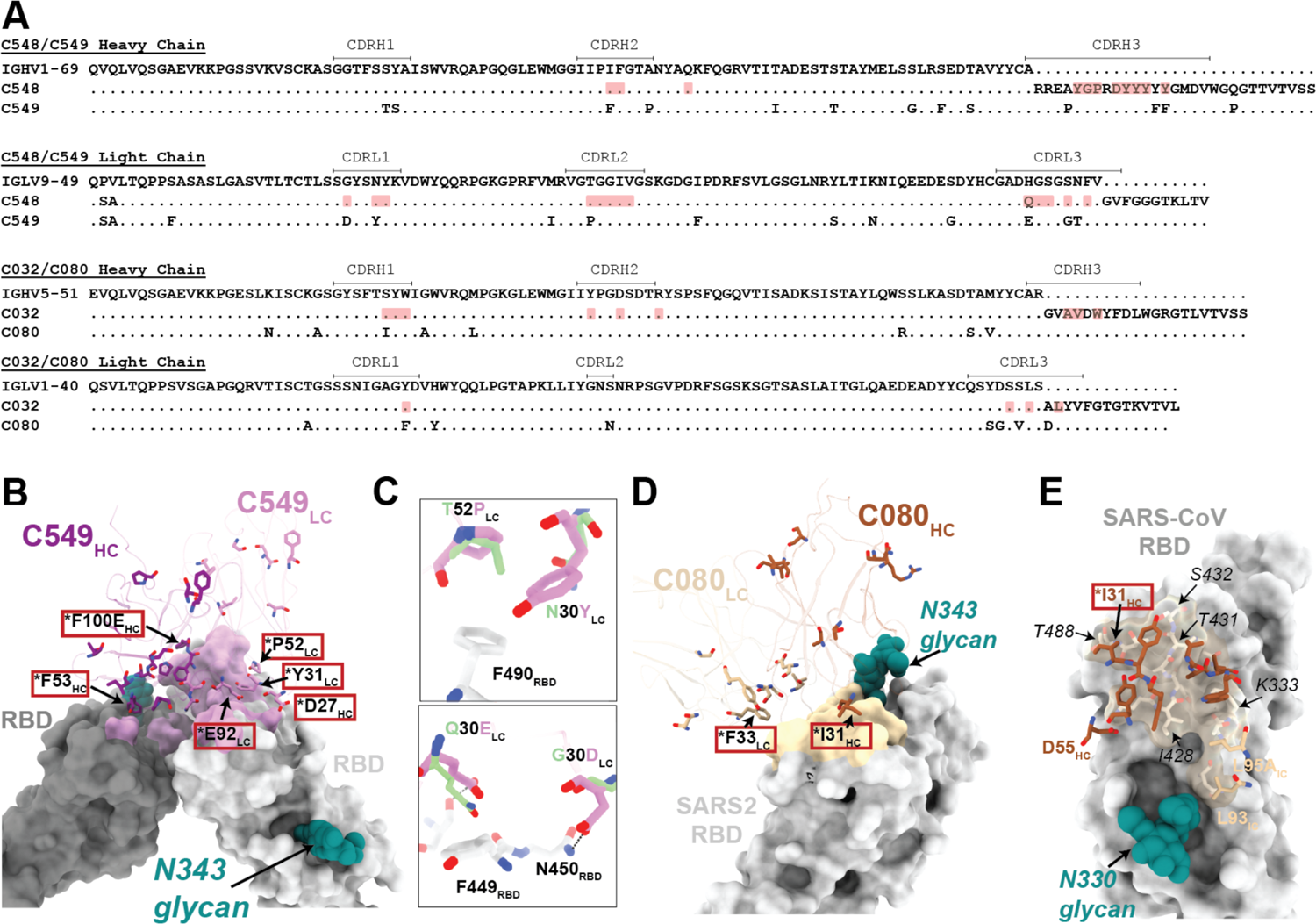
Class 2 and 3 antibody sequence alignments and homology models, related to Figure 7. (A) Sequence alignment between heavy and light chains of C548/C549 and C032/C080 antibody pairs relative to inferred germline sequences. Paratope residues highlighted in red. (B) Homology model of C549-RBD complex. Antibody somatic mutations are shown as sticks. Residues changed by somatic hypermutation at the predicted RBD interface are indicated by an asterisk and enclosed in a red box. (C) Predicted interactions between RBD (light gray) and C549 homology model LC residues (violet). C548 residues (light green) are shown. (D) Homology model of the C080-RBD complex. Antibody somatic mutations are shown as sticks. Residues changed by somatic hypermutation at the predicted RBD interface are indicated by an asterisk and enclosed in a red box. (E) Homology model of the C080-SARS-CoV RBD complex. Predicted RBD epitope and Fab paratope are shown as colored surface and sticks, respectively. Residues changed by somatic hypermutation at the predicted RBD interface are indicated by an asterisk and enclosed in a red box. Sequence differences in SARS-RBD relative to SARS-CoV-2 RBD are indicated with italics.

### Supplemental Tables S1-S3

**Table Sl.**
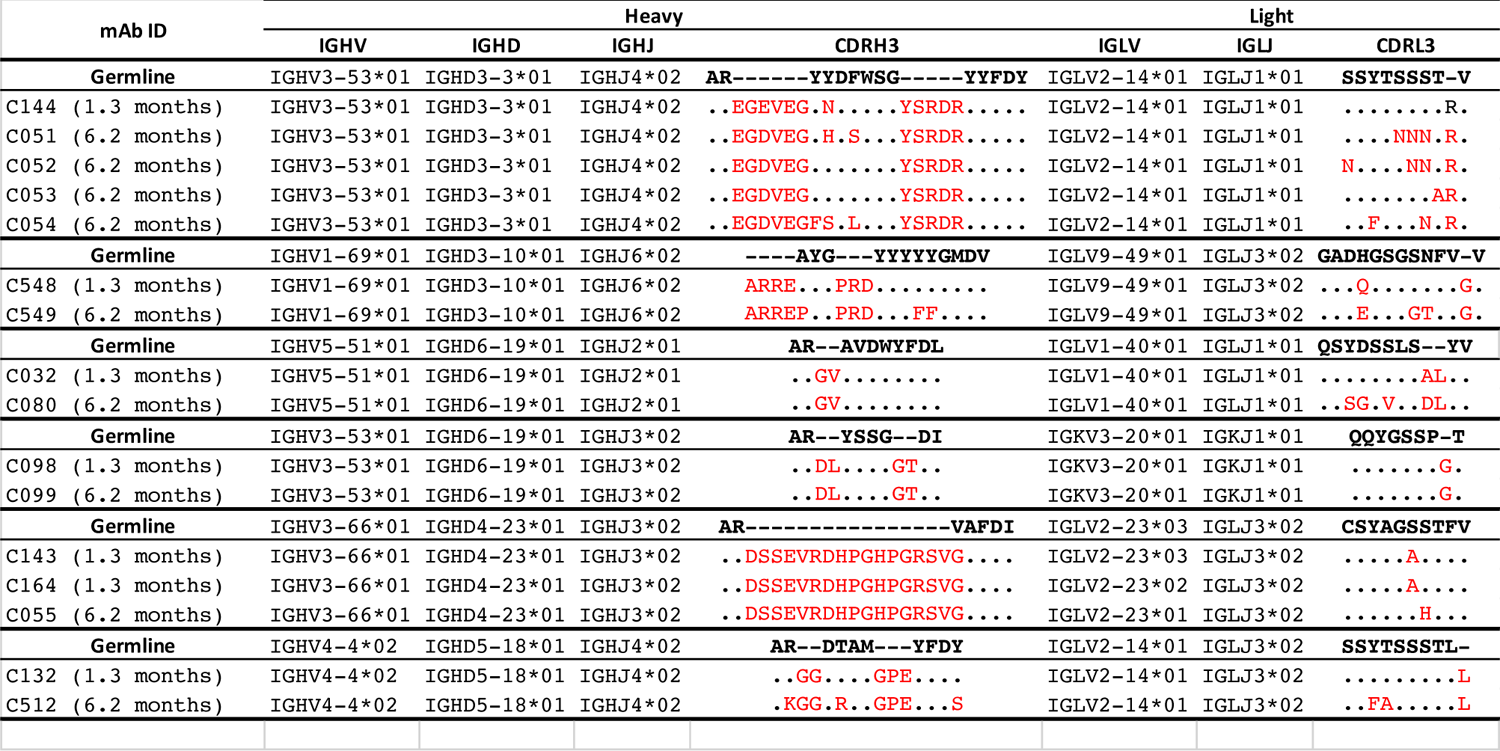
Clonally related antibody lineages in this study

**Table 82.**
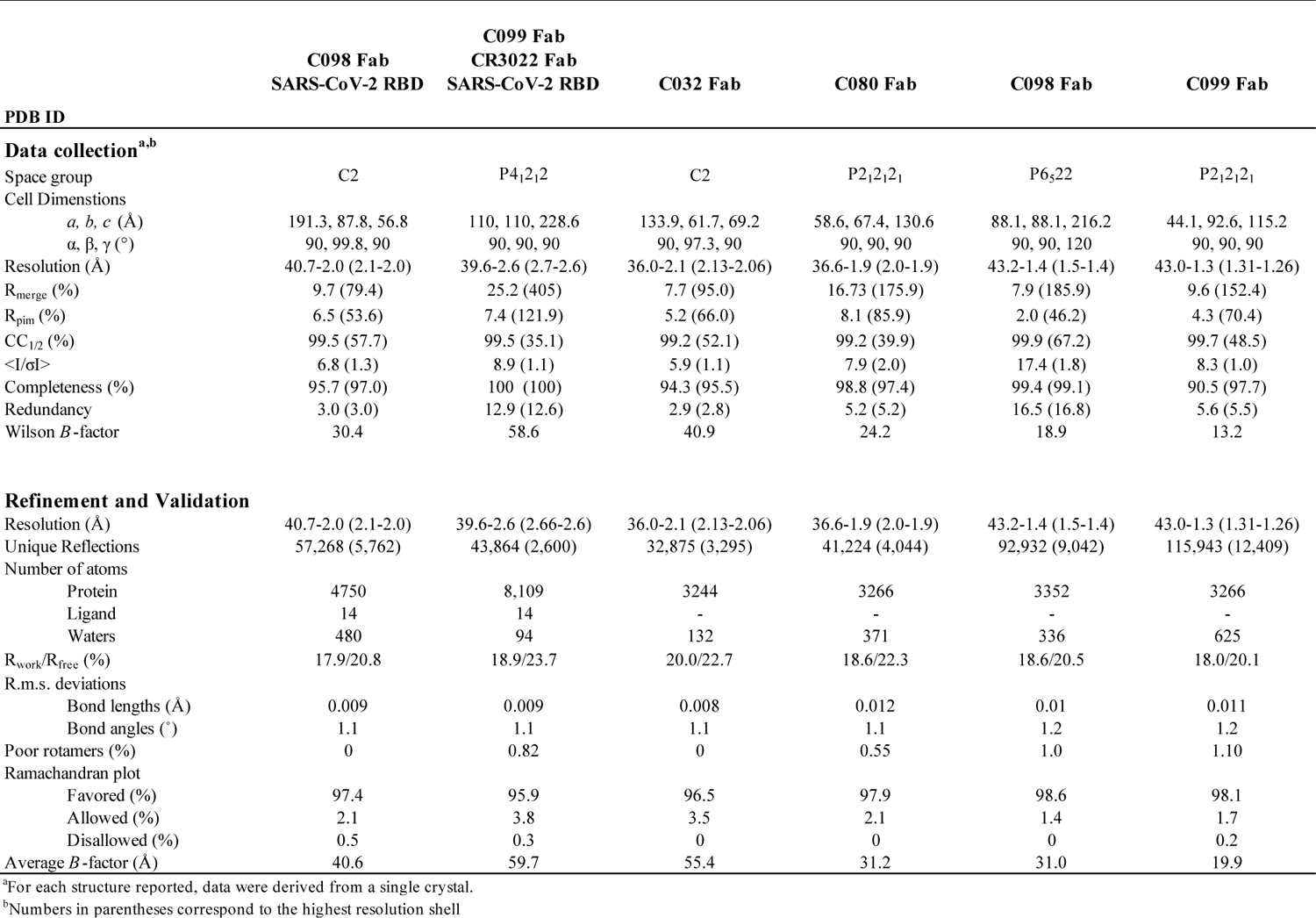
X-ray data collection and refinement statistics.

**Table S3.** ct-yo-EM data collection and refinement statistics.

|  | **C032** | **C051** | **C548** |
| --- | --- | --- | --- |
|  | **SARS-CoV-2 S6P** | **SARS-CoV-2 S 6P** | **SARS-CoV-2 S 6P** |
| **PDB** |  |  |  |
| **EMD** |  |  |  |
| **Data collection conditions** |  |  |  |
| Microscope | Talos Arctica | Talos Arctica | Talos Arctica |
| Camera | Gatan K3 Summit | Gatan K3 Summit | Gatan K3 Summit |
| Magnification | 45,000x | 45,000x | 45,000x |
| Voltage (kV) | 200 | 200 | 200 |
| Recording mode | counting | counting | counting |
| Dose rate (e<sup>-</sup>/pixel/s) | 13.5 | 13.5 | 13.5 |
| Electron dose (e<sup>-</sup>/Å<sup>2</sup>) | 60 | 60 | 60 |
| Defocus range (μm) | 0.7 – 2.0 | 0.7 – 2.0 | 0.7 – 2.0 |
| Pixel size (Å) | 0.8689 | 0.8689 | 0.8689 |
| Micrographs collected | 3,480 | 3,402 | 1959 |
| Micrographs used | 2,683 | 2,927 | 1687 |
| Total extracted particles | 844,544 | 390,630 | 554,852 |
| Refined particles | 192,286 | 134,506 | 94,255 |
| Symmetry imposed | C1 | C3 | C3 |
| Nominal Map Resolution (Å) |  |  |  |
| FSC 0.143 (unmasked/masked) | 5.4/3.4 | 4.9/3.5 | 4.5/3.5 |
| FSC 0.143 local (unmasked/masked) | 6.7/4.4 | 5.8/4.1 | N/A |
| **Refinement and Validation** |  |  |  |
| Initial model used | 6XKL | 7K90 | 7K43 |
| Number of atoms |  |  |  |
| Protein | 25,866 | 28,401 | 28136 |
| Ligand | 56 | 462 | 574 |
| MapCC (global/local) | 0.70/0.69 | 0.80/0.77 | 0.77/0.76 |
| Map sharpening B-factor | 69.3 | 64.5 | 110.1 |
| R.m.s. deviations |  |  |  |
| Bond lengths (Å) | 0.004 | 0.008 | 0.003 |
| Bond angles (°) | 0.64 | 1.1 | 0.52 |
| MolProbity score | 2.18 | 2.29 | 2.48 |
| Clashscore (all atom) | 16.3 | 16.4 | 18.26 |
| Poor rotamers (%) | 0 | 0.2 | 0.38 |
| Ramachandran plot |  |  |  |
| Favored (%) | 92.4 | 93.8 | 97 |
| Allowed (%) | 7.5 | 6.9 | 2.7 |
| Disallowed (%) | 0 | 0.3 | 0.3 |

